# Structure and Protein-Protein Interactions of Ice Nucleation Proteins Drive Their Activity

**DOI:** 10.1101/2022.01.21.477219

**Authors:** Susan Hartmann, Meilee Ling, Lasse S.A. Dreyer, Assaf Zipori, Kai Finster, Sarah Grawe, Lasse Z. Jensen, Stella Borck, Naama Reicher, Taner Drace, Dennis Niedermeier, Nykola C. Jones, Søren V. Hoffmann, Heike Wex, Yinon Rudich, Thomas Boesen, Tina Šantl-Temkiv

**Affiliations:** Institute for Tropospheric Research, Permoserstraße 15, Leipzig, 04318, Germany; Department of Biology, Microbiology Section, Aarhus University, Ny Munkegade 114-116, Aarhus 8000, Denmark; Department of Physics and Astronomy, Stellar Astrophysics Centre, Aarhus University, Ny Munkegade 120, Aarhus 8000, Denmark; Department of Molecular Biology and Genetics, Section for Protein Science, Aarhus University, Gustav Wieds Vej 10 C, Aarhus 8000, Denmark; Department of Earth and Planetary Sciences, Weizmann Institute of Science, Rehovot 76100, Israel; ISA, Department of Physics and Astronomy, Aarhus University, Ny Munkegade 120, Aarhus 8000, Denmark; Interdisciplinary Nanoscience Center, Aarhus University, Gustav Wieds Vej 14, Aarhus 8000, Denmark

**Author notes:** Corresponding Authors: Thomas Boesen, Gustav Wieds Vej 10 C, Aarhus 8000, Denmark, ORCID: 0000-0002-5633-6844; Tina Šantl-Temkiv, Ny Munkegade 114-116, Aarhus 8000, Denmark, ORCID: 0000-0002-3446-5813. These authors contributed equally (Susan Hartmann, Meilee Ling, Lasse S.A. Dreyer).

## Abstract

Microbially-produced ice nucleating proteins (INpro) are unique molecular structures with the highest known catalytic efficiency for ice formation. Their critical role in rain formation and frost damage of crops together with their diverse commercial applications warrant an in-depth understanding of their inherent ice nucleation mechanism. We used the machine-learning based software AlphaFold to develop the first *ab initio* structural model of a bacterial INpro which is a novel beta-helix structure consisting of repeated stacks of two beta strands connected by two sharp turns. Using the synchrotron radiation circular dichroism, we validated the β-strand content of the model. Combining functional studies of purified recombinant INpro, electron microscopy and modeling, we further demonstrate that the formation of dimers and higher-order oligomers is key to INpro activity. This work presents a major advance in understanding the molecular foundation for bacterial ice-nucleation activity and the basis for investigating the mechanistic role of INpro-induced ice formation in the atmosphere, and for commercial design and production of ice-nucleating particles for industrial applications.

## 1. Introduction

Microbially-produced ice nucleating proteins (INpro) are the most efficient catalysts for ice formation in nature ^1^. They are ubiquitous in terrestrial, marine and atmospheric habitats as well as on microbial, plant, fungal and animal surfaces and have been shown to maintain their activity for at least 30,000 years ^2–4^. INpro are unique in that they can initiate freezing just below 0°C, i.e. close to the melting point of ice ^1, 5^. While pure water freezes close to -38 °C, ice frequently forms close to 0°C ^6^, which underlines the enormous environmental relevance of INpro, including extensive frost damage of crops and impact on cloud radiative properties and lifetime ^7–11^, as well as their commercial applications, including artificial snow production, cloud seeding, crop protection and food preservation ^12–15^. Currently known microbial sources of INpro are plant-associated bacteria of the genera *Pseudomonas*, *Pantoea* and *Xanthomonas* ^16, 17^, soil fungi of the genera *Fusarium* ^18^ and *Mortierella* ^19^ and several species of terrestrial and marine Cyanobacteria and microalgae ^20, 21^. Despite the diverse sources of INpro, bacterial INpro are the only verified INpro with known amino acid sequences ^22^ and are therefore widely used as molecular models for INpro. However, for decades bacterial INpro have eluded structural analysis due to their large size and their association with cell membranes. Very recently, a new version of the structure prediction tool AlphaFold has been released as the first computational method that can predict atomic three-dimensional protein structures with high confidence, as validated by comparison to experimental structures. The machine learning (ML) based software is a major improvement in comparison to existing structure prediction algorithms, and is a powerful tool to be used when experimental structures remain elusive ^23^.

Bacterial INpro are membrane-associated proteins with molecular weights in the range of 150-180kDa^22^. INpro are believed to belong to a family of proteins that contain tandemly repeated, elongated, open structures with a theoretically unlimited number of repeats ^24^. The central repeat domain (CRD), which is composed of a variable number of 16-amino-acid tandem repeats with the consensus sequence “GYGSTxTAxxxSxL[T/I]A”, constitutes the largest part of the INpro molecule ^25, 26^. Generally, the functional advantage of repeats has been associated with the enlargement of the available binding surface ^27^, which is involved in ligand binding ^24^. In the case of INpro molecules, the active-surface area has been suggested to facilitate binding and ordering of water molecules, and thereby the protein would act as a template for the formation of an ice nucleation embryo and thus induce freezing in the whole volume of water ^28, 29^.

Bacterial INpro show two classes of ice nucleation activity that are characterized by distinct nucleation temperatures ^30^. The INpro class C, which nucleate ice between −7 °C and −10 °C, are frequently observed while INpro class A, which nucleate ice between −2 °C and −5°C, are rare. While there is a general consensus that the temperature at which INpro nucleate ice correlates with INpro oligomer size ^17, 31–33^ and INpro class A has recently been shown to depend on electrostatic interactions ^34^, the experimental evidence showing assembly of INpro into oligomers and the underlying mechanism that correlates oligomeric state and ice nucleation activity has yet to be provided.

Despite their multifaceted environmental impacts, only a limited number of modellingand experimental studies on the structures and interactions of bacterial INpro has been carried out ^17, 25, 35–39^. For example, based on a homology modelling study, it has been suggested that each of the 16 amino acids repeats forms a right-handed β-helix with a triangular base and three β-sheet faces, which aligns the threonine-x-threonine motifs (TxT motif) along one face of the β-helix and the serine-x-leucine-threonine/isoleucine (SxL[T/I] motif) along the other face, with each motif forming a short β-sheet. Ac-cording to this model, the two β-sheets defined by these faces are the sites where ice nucleation takes place. The authors furthermore hypothesized that two monomers can dimerize along the conserved tyrosine ladder located in the third β-sheet face of the β-helix. This dimerization was suggested to connect the TxT motif of one monomer with the SxL[T/I] motif of the other monomer, thus extending the surface for ice-nucleation ^25^. Founded on the outcome of molecular dynamics simulations using this INpro model ^25^, Hudait et al. (2018) suggested that the TxT and SxL[T/I] β-sheets have comparable ice nucleation properties and promote ice formation through an anchored clathrate and ice-like motifs, respectively ^36^. Structural modelling of the CRD has so far been derived from the initial homology model proposed by Garnham *et al*. ^25^.

Using interface-specific sum frequency generation (SFG) spectroscopy in combination with modelling, Pandey et al. (2016) proposed that the extended domain of aligned TxT motifs enhances ice-nucleation through a hydrophilic-hydrophobic pattern ^35^. Recently, by combining SFG with two-dimensional infrared spectroscopy, Roeters et al. (2021) provided support for the β-helical INpro structure and showed that the protein imposes order on adjacent water molecules, in particular at low temperature ^40^. Ling et al. (2018) demonstrated that INpro molecules maintain ice-nucleation capacity despite a 4-fold reduction in the molecular size, and that the ice nucleation temperature decreased as a function of a reduced number of repeats ^17^. In spite of these pioneering studies, we are still in need of experimentally verified insights into the link between the structural and functional properties of the INpro molecules.

In this paper we report on the first experimentally validated model of a bacterial INpro. We reached our goal by combining *ab initio* modeling of the INpro 3D molecular structure with synchrotron radiation circular dichroism (SRCD) and transmission electron microscopy (TEM) analysis as well as ice nucleation assays and modeling of ice cluster formation based on Classical Nucleation Theory (CNT). Combing these complementing methods, we were able to considerably advance our knowledge of the effect of INpro structure and molecule interaction on their ice nucleation activity.

## 2. Results and Discussion

### 2.1. The INpro repeats form a β-helix structure

Combining ML-based structure prediction of AlphaFold ^23^ with synchrotron radiation circular dichroism data, we propose a new model for the structure based on the initial 16 repeats of the INpro CRD from *Pseudomonas syringae* R10.79 ^41^. The model was based on the initial 16 N-terminal repeats, as this repeat section displays the greatest sequence conservation across repeats. The proposed model (Figure 1A, Figure S1) consists of two β-strands and connected by two sharp turns. The two β-strands form opposing parallel β-sheets. By the interactions between neighboring repeats a β-helix structure is formed with an unusual polar core between the β-sheets. The two strands have a limited hydrophobic core consisting of inward facing leucine and alanine residues. Sharp turns are facilitated by the conserved glycine residues in the repeat sequence, and are stabilized by two internal serine ladders – one in each turn (Figure 1B). This differentiates our model from the earlier homology model proposed by Garnham et al. ^25^, which produces a wider loop-like structure in the tyrosine turn forming a third β-strand with no internal serine ladder. Despite these differences, both models share the solvent-exposed tyrosine ladder on one of the turns. The tyrosine residues are fully conserved in the first 59 repeats of the CRD. One β-sheet is decorated with the proposed TxT ice-nucleating motif while the other β-sheet carries the proposed SxL[T/I] ice-nucleating motif. Both β-strands in the repeat form a locally flat surface, but the β-helix itself has a rotation along the longitudinal axis of approx. 45 degrees from N- to C-terminal. The two putative ice-nucleating motifs are pointing in opposite directions relative to the longitudinal β-helix axis. The consensus sequence of the CRD is shown in Figure 1C with annotations of predicted structural features. The model of the complete INpro having 67 repeats and short N- and C-terminal domains is presented in Figure S2 and exhibits very similar features as the model of the initial 16 N-terminal repeats.

**Figure 1:**
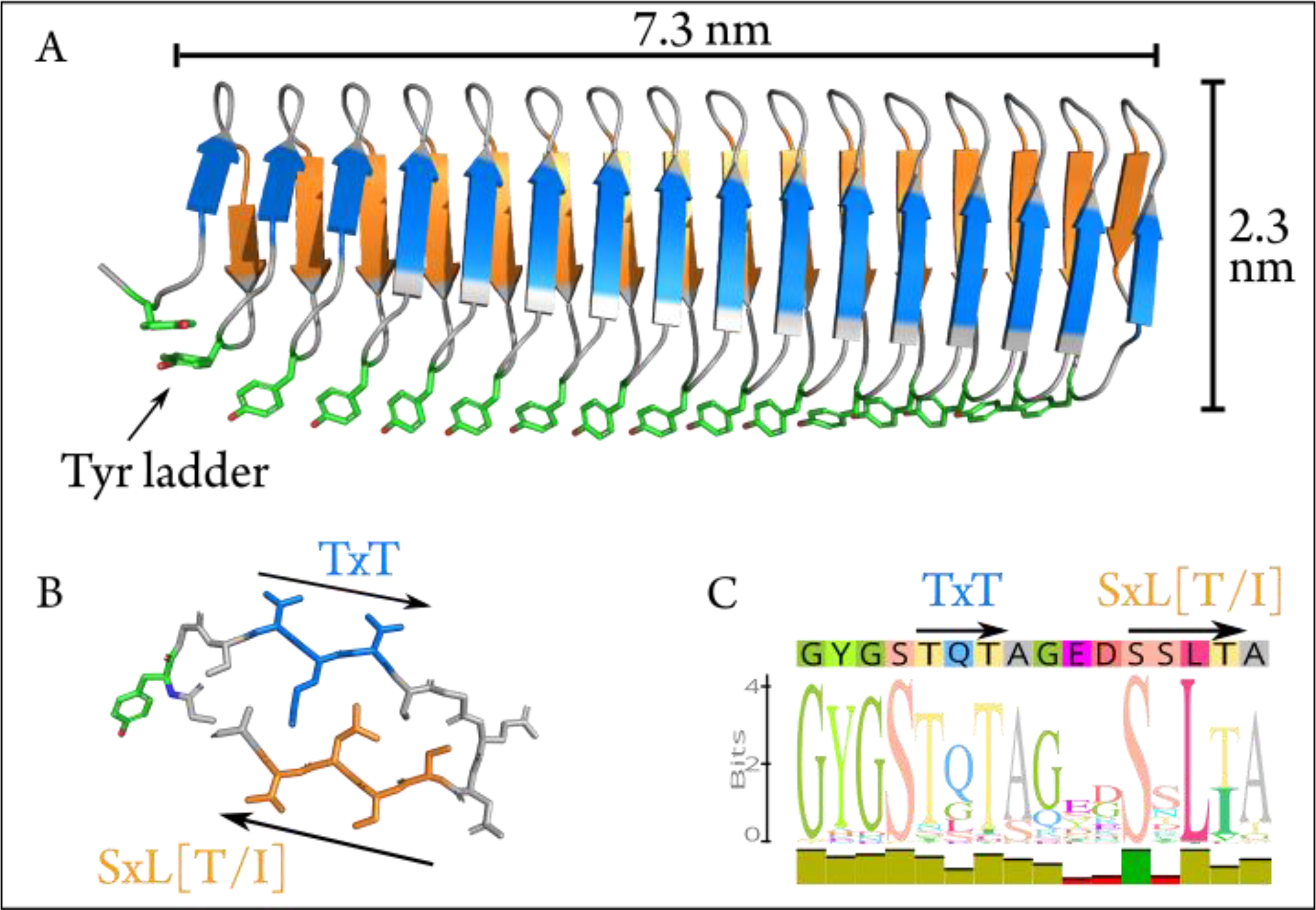
*Ab initio* model of the first 16 repeats of the INpro CRD. **(A)** The *ab initio* model predicted using machine-learning-based algorithms in AlphaFold. The model consists of a β-helix with one extended β-sheet on each side. The β-strands have a rotation along the longitudinal axis of approximately 40 degrees when comparing Nto C-terminal. The highly conserved tyrosine ladder (annotated) is solvent-exposed along the side (shown with stick-representation). **(B)** Stick representation of repeat 13 which displays the best match to the consensus sequence (shown in **C**). The two putative ice-nucleation active sites are annotated as TxT and SxL[T/I], respectively. **(C)** The consensus sequence of the INpro CRD. The predicted structural features are annotated above the sequence corresponding to the color scheme in A and B.

Using the INpro sequence of *P. syringae* R10.79 ^41^ with a total number of 67 repeats, we designed and produced four recombinant INpro constructs with 9, 16, 28 and 67 tandem repeats in the CRD, i.e. INpro-9R, INpro-16R, INpro-28R and INpro-67R, respectively (Figure 2A, Figure S3, Figure S4, Table S1). The constructs were expressed in *E. coli* and purified (Table S2).

**Figure 2:**
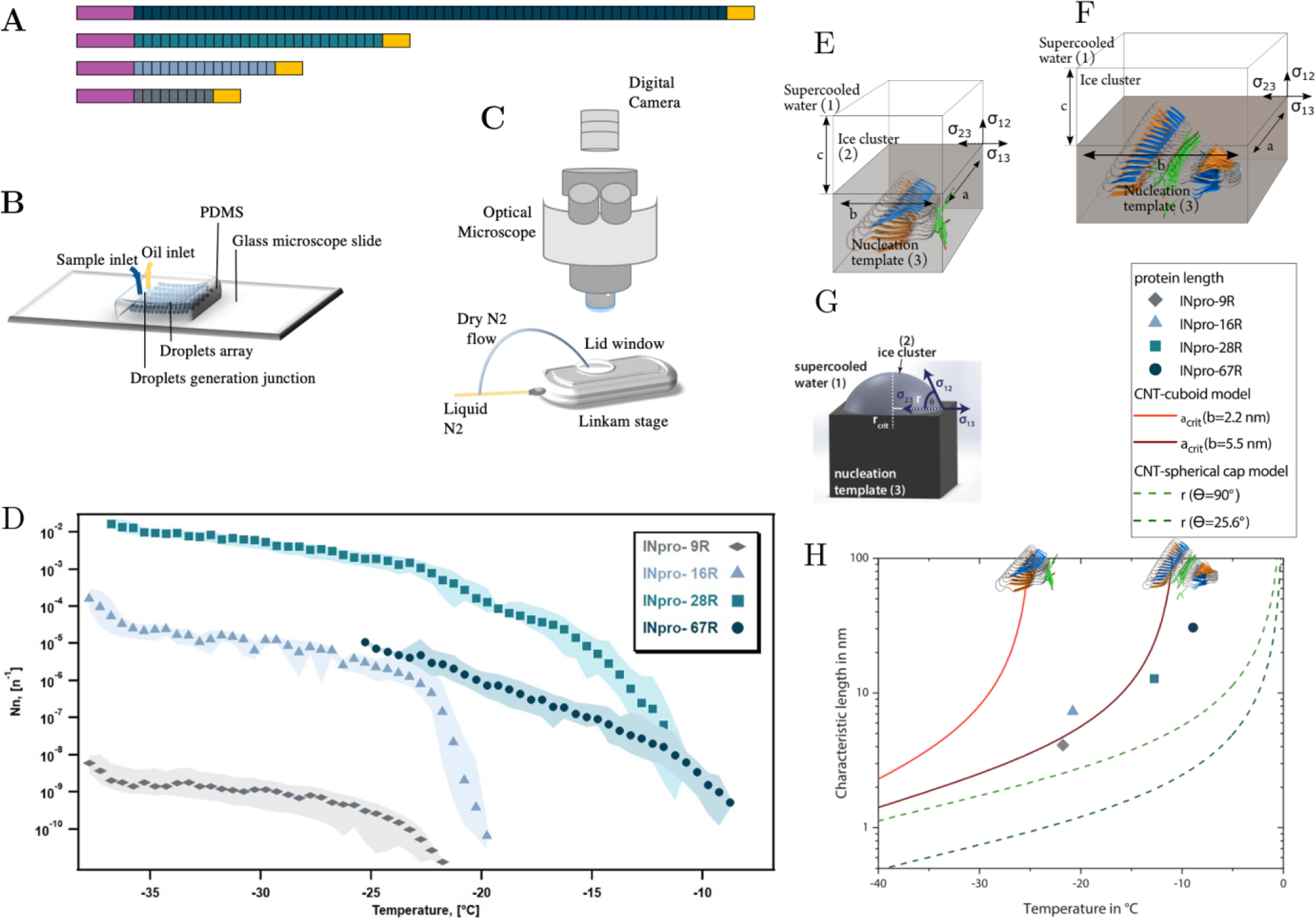
The central repeat region (CRD) is an elongated template for ice nucleation. **(A)** Four INpro constructs were produced to investigate the role of CRD size for the nucleation activity. Purple signifies the N-terminal domain and the yellow signifies the C-terminal domain. The colors of the CRD are used to differentiate the recombinant INpro constructs. **(B)** A schematic drawing of WISDOM instrument that was employed to investigate the ice-nucleation activity of recombinant INpro constructs. Patterned PDMS (polydimethylsiloxane) attached to a microscope glass slide was used to form the microfluidic device. Oil and sample flows meet in a junction where nanoliter monodisperse droplets are generated and subsequently arranged in a microfluidic chamber array. **(C)** WISDOM freezing experiments. The microfluidic device is placed in a cooling stage (Linkam) coupled to an optical microscope and a CCD camera. During the experiment a flow of dry N_2_ was used to prevent condensation on the outer window. **(D)** Ice nucleation spectra of INpro constructs expressed as ice nucleation site density per molecule (*N*_*n*_) as a function of temperature. Each color represents a protein of different length. More detailed figures showing ice nucleation spectra of each protein construct can be found in the SI (Figure S6). The quality with which the steepness of the increase was captured depends on the concentration of the purified INpro molecules in each sample. **(E, F, G)** A sketch of the ice nucleation template (3) and the ice cluster (2) in supercooled water (1) for cuboid geometry proposed based on the *ab initio* β- helical INpro model **(E and F)** and spherical cap geometry previously used to model INpro activity **(G)**. The interfaces are defined via the respective interfacial free energies σ_ij_ of the adjoining phases. The cuboid is determined by three axes (a, b, c), whereas the spherical cap can be described by the critical radius r_crit_ and the contact angle θ. **(H)** Comparison of the relationship between the characteristic length (a or r in Figures 2E and 2F) and the temperature predicted from the cuboid and spherical cap models according to CNT and the experimental measurements of characteristic nucleation temperatures and lengths of INpro dimers determined from the *ab initio* β-helical model of the INpro ^25^.

To validate the new INpro repeat model, we set out to obtain the first experimental secondary structure data of individual repeats based on recombinant proteins having both the N- and C-terminal domains flanking the CRD domain. Purified INpro-9R and INpro-16R samples were analysed using SRCD spectroscopy and data for multiple concentrations of each sample were deconvoluted using the CDSSTR analysis programme and SMP180 reference datasets at the DichroWeb portal ^42, 43^, thereby confirming consistency of the derived secondary structure content (Figure S5). The secondary structure content for the 7 repeats representing the difference between the two constructs was calculated to be ∼35% β-strand, the remaining secondary structure being turns and coil structures. This agrees better with the lower β- strand content (∼50%) of our *ab initio* model of a two-β-strand β-helix structure than the three-β-strand β-helix model proposed by Garnham et al. (2011) that has a β-strand content of ∼69% ^25^. The difference between a secondary content of 35% and 50% corresponds to each of the strands being one amino acid shorter and these amino acids would then be part of the loop/coil structures. It is possible that the reference data sets (based on known structures) used for deconvolution of the circular dichroism data resulted in the assignment of these amino acids to coil/loops instead of β-strands in the unusual INpro β-helix structure.

### 2.2. Dimerization on INpro is a prerequisite to class-C ice-nucleation activity

We used purified INpro-9R, INpro-16R, INpro-28R and INpro-67R to investigate the role of the CRD’s repeat number for the ice nucleation activity of INpro (Figure 2A, Figure S3, Figure S4, Table S1, Table S2) using the WISDOM setup (Figure 2B and C) ^44^. This setup utilizes nanoliter size droplets and allows ice nucleation measurements down to – 38°C, the temperature of homogeneous freezing of water. Figure 2D (and Figure S6) depicts ice nucleation site density per molecule (*N*_*n*_) as a function of temperature, which we termed the ice nucleation spectrum, for the four different INpro constructs. The shift from a steep to a gentle increase in the ice nucleation spectrum was defined as the knee point of the curve. This can best be seen for INpro-16R that has a knee point just below -20°C (Figure 2D, Figure S7, Table S3). The other constructs show comparable features and this allowed us to identify the knee points for all ice nucleation spectra (Figure S7). The steepness of the ice nucleation spectrum depends on the homogeneity of the INpro molecules present in a sample. Therefore, the steep slopes could either be associated with the activity of functional INpro monomers or with the presence of homogeneous oligomers. We used the knee point to define T_char,50_ , i.e. the characteristic nucleation temperature of homogeneous INpro at 50% of the concentration ovserved at the knee point (Table S3). The calculated T_char,50_ were -23.7 °C, -21.7 °C, -15.8 °C and -11.2 °C for INpro-9R, INpro-16R, INpro-28R and INpro-67R, respectively. Our results clearly show that ice nucleation activity scales with the number of repeats (Figure 2D) but not in a linear fashion. Thus, the relative increase in T_char,50_ as a function of repeat number was less pronounced between INpro-28R and INPro-67R than between INpro-9R and INpro-16R or INpro-16R and INpro-28R (Figure 2D).

In previous studies, it has been hypothesized that the size of ice nucleation active molecules plays a major role in determining their characteristic nucleation temperature ^17, 29, 31, 45, 46^. To test this hypothesis, we derived the transverse and longitudinal axes of the constructs using our structural INpro model. We estimated that while the transverse axis of the INpro monomers is 2.3 nm in all cases, the longitudinal axis of the CRD is 4.1 nm in the INpro-9R monomer, 7.3 nm in the INpro-16R monomer, 12.8 nm in the INpro-28R monomer and 30.6 nm in the INpro-67R monomer (Table S3). We based the rectangular template shape catalyzing cuboidal ice cluster formation on the slim, elongated shape of the INpro molecular model (Figure 2E). The rectangular template is described by the length of its longitudinal and transverse axis. During the formation of ice clusters in supercooled water, energy is released as the volume of the ice cluster increases, while energy is needed to form the interfaces between the ice cluster and the surrounding liquid. A combination of the volumeand the surface effect defines the critical energy barrier for the ice cluster formation. Finally, one of the many continuously forming and melting ice clusters overcomes the critical energy barrier and initiates freezing of the whole supercooled water volume. It is energetically favorable that this process takes place at the surface of the nucleation template. From the critical energy barrier, we derive the respective critical length of the longitudinal axis that forms the critical ice cluster (a_crit_, Figure 2E), assuming a constant transverse axis (b).

Assuming a cuboidal geometry of the ice cluster, we obtain a better agreement with data from direct measurements of INpro constructs compared to the standard CNT model, which assumes a spherical cap geometry (Figure 2E-H). Based on the cuboid model, a single INpro molecule with a width of 2.2 nm is a very poor ice nucleator. Our experimental data correlate well with the double transverse axis length (baxis) of 5.5 nm (Figure 2F and H).

As the combination of the CNT model and experimental data suggests that the ice-nucleating surface is wider than an INpro monomer, we performed computational docking based on rigid-body algorithms to propose a homodimer structure of the initial 16 repeats of the INpro CRD (presented in section 2.1). The proposed dimer structure is presented in Figures 3A and B. The monomers align along the highly conserved tyrosine ladder. This is in good agreement with earlier molecular dynamics simulations ^25^. The dimer structure is parallel, meaning that the two monomers are oriented N- to N-terminal, resulting in a surface where the TxT motif of one monomer aligns with the SxL[T/I] motif on the other monomer. During multiple docking runs with varying parameters the parallel dimer was consistently selected over the anti-parallel by the docking algorithm. The transverse axis of the dimer is 5.5 nm (Figure 3A, TxT and SxL[T/I] marked in blue and orange, respectively), which corresponds very well with the experimentally-measured characteristic nucleation temperatures (Figure 2H). Interestingly, the longitudinal rotation of the β-helix present in each monomer leads to a less flat surface of the dimer when docked along the tyrosine ladder. Instead, the surface adopts a saddle-like structure. This is clearly visible in Figure 3B. As the INpro is bound to the outer membrane, the topology of a dimer and higher-order oligomers will likely be influenced by *in situ* conditions, e.g. membrane curvature assuming that the INpro oligomer is lying flat on the cell surface and interacts with the outer membrane. Nevertheless, the dimensions of a dimer used in the CNT modelling are in good agreement with the experimental data on protein activity (Figure 2H), strongly indicating that the INpro molecules have to form dimers to be exhibit observed ice-nucleation activity. We therefore suggest that dimers are responsible for INpro class C ice nucleation.

**Figure 3:**
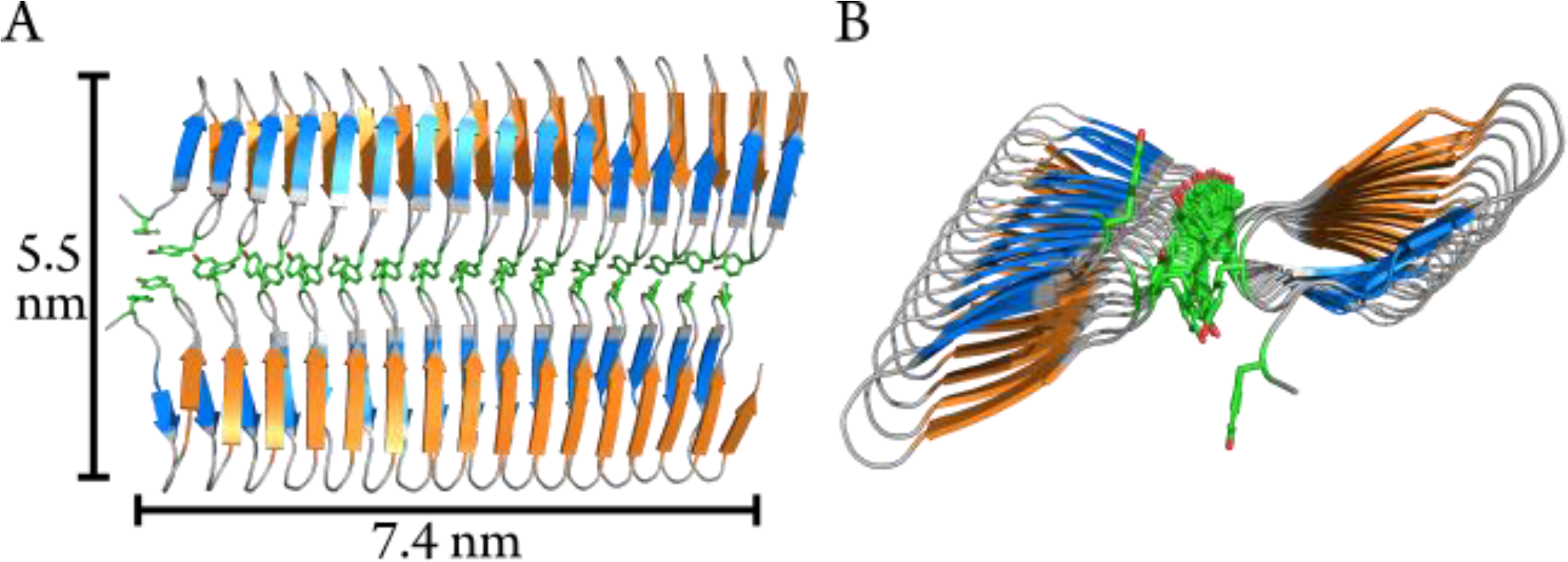
Modelled homodimer structure of the initial 16-repeats of the INpro CRD domain. (**A**) The proposed homodimer structure of the INpro CRD. The tyrosine ladder comprises the dimerization interface. The monomers are parallel, and the TxT motif of one monomer is aligned with the SxL[T/I] motif of another monomer (blue and orange, respectively). The tyrosine ladder forms the dimerization interface. Approximate dimensions of the dimer surface are indicated. **(B)** End-view of the dimer model. The longitudinal rotation in the β-helix causes the dimer to form a saddle-like surface.

### 2.3. Class-A ice-nucleation activity is linked to higher-order INpro oligomers

Based on the cuboid model used to obtain data presented in Figure 2H, we conclude that the effect of additional amino-acid repeats in the CRD on the ice nucleation temperature is high when the number of repeats in the CRD is low and decreases as the number of repeats in the CRD is higher. This is due to the fact that the characteristic nucleation temperature, at a certain transverse axis length, is approached asymptotically as a function of increasing number of repeats. This led us to conclude that oligomerization of the INpro dimers into higher order filamentous structures ^39, 47^ raises the characteristic ice nucleation temperature more effectively than increasing the length of a single INpro molecule by adding more repeat modules.

Using the LINA setup (Figure 4A and 4B) ^48^, which measures ice nucleation using microliter size droplets, we studied whether the less abundant class A INpro activity only occurs in the presence of cells and membrane lipids. Investigating *E. coli* cells that express INpro-67R, we confirmed the presence of two INpro classes (Figure 4C): Class A INpro with T_char,50_ of -3.9°C were ∼2 orders of magnitude less abundant than Class C INpro with T_char,knee_ of -7.6°C (Table S3). Our findings are in agreement with recent high-resolution ice-nucleation studies of the same INpro type ^49, 50^ (Figure 4C). The steep slopes observed for the two INpro classes are best explained by two populations of homogenous INpro, while the plateau between the slopes results from the fact that all Class A INpro are already activated at a certain temperature (T_char,50_, Table S3). Oligomeric structures of INpro assembling on the outer cell membrane were proposed to act as class A INpro ^28, 31, 33, 51^. We observed that at high INpro concentration, purified INpro-67R also contain class A INpro (Figure 4C, Table S3). This observation confirmed that the activity of class A INpro does not depend on the presence of bacterial cells or membrane lipids as previously thought ^52^. A result that is further supported by the fact that solutions of the purified INpro-16R constructs contain molecular structures with a high level of activity (T_char,50_ of -9.3°C, see at > -10°C in Figure 4C, not included in Table S3) and structures with a low level of activity (T_char,50_ of -22.2°C, see < - 15°C in Figure 4C, Table S3). The two structures of INpro-16R were found at a similar concentration ratio as those observed for the classes in purified INpro-67R. Thus, the two structures of INpro-16R are very likely homologous to class A and class C INpro-67R. Based on these results, we hypothesize that while class C INpro are composed of INpro dimers that induce ice nucleation in the low temperature range, class A INpro are composed of higher order INpro oligomers that induce ice nucleation in the high temperature range. Both INpro classes form in solution from purified proteins in the absence of cells and membranes. We observed a large difference in concentration between the two classes of INpro, which could be attributed either to (i) a higher concentration of INpro dimers compared to INpro oligomers, or to (ii) a concentration-dependent self-assembly of INpro oligomers from the INpro dimers or to (iii) a combination of (i) and (ii) considering a reversible oligomerization process.

**Figure 4:**
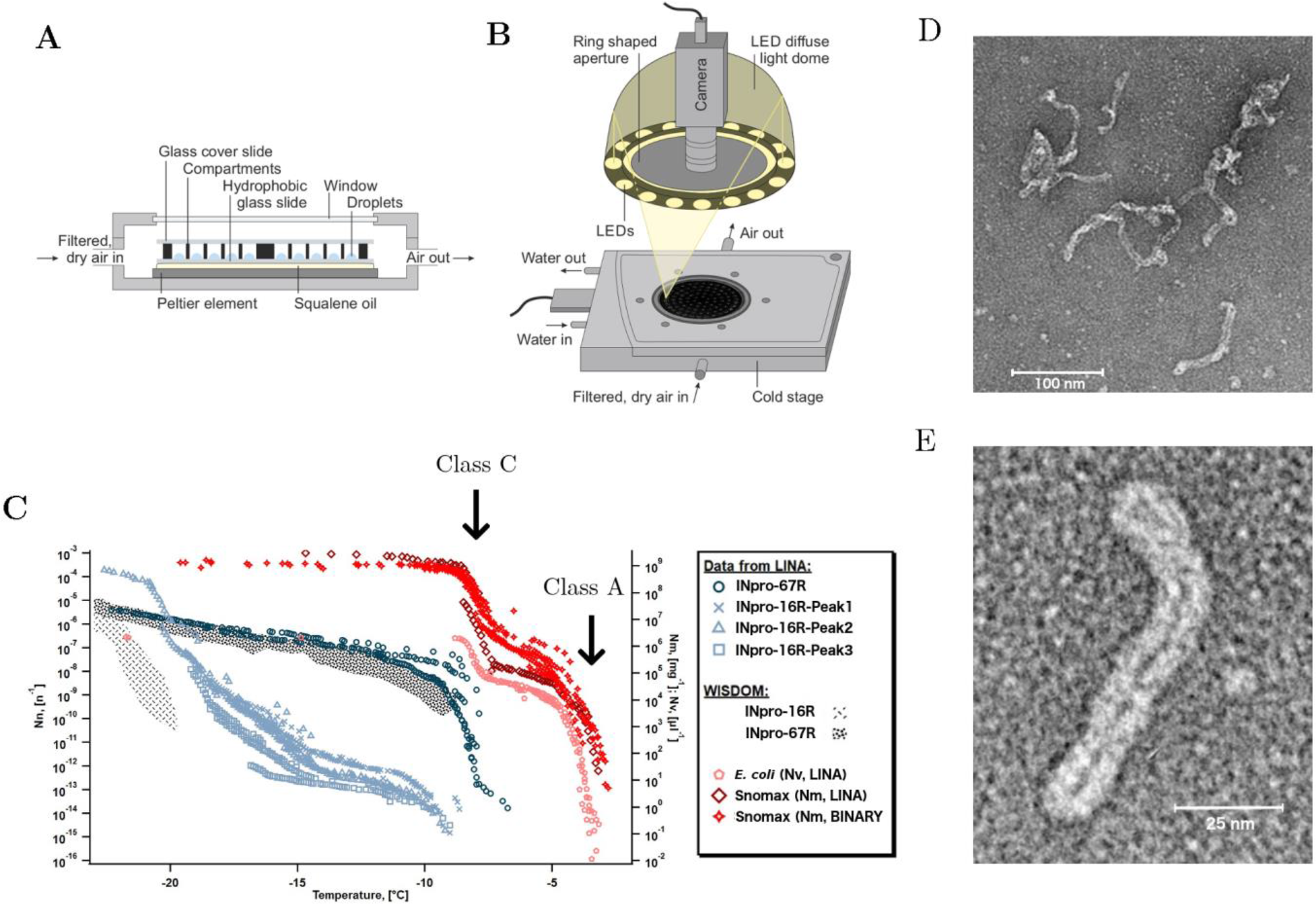
Two types of INpro oligomers are responsible for the ice-nucleation activity of two INpro classes. **(A and B)** A schematic drawing of the LINA setup used to investigate the ice nucleation activity of the two INpro classes. **(C)** Ice nucleation activity of *E. coli* expressing INpro-67R as well as of purified INpro-67R and INpro-16R proteins measured by the LINA setup. Presumed class A and class C characteristics and structures are indicated for *E. coli* and Snomax® ice nucleation spectra, but are also seen in INpro-67R and INpro-16R spectra. Snomax® data were obtained from previous studies ^49, 50^. Data from WISDOM ice-nucleation measurements of INpro-16R and INpro-67R shown for comparison. **(D and E)** Representative negative stain TEM images of the INpro-67R showing highly oligomerized, filamentous structures.

During purification, all proteins (INpro-9R, INpro-16R, INpro-28R and INpro-67R) showed multiple peaks in size exclusion chromatography (SEC), e.g. Figure S8A. The presence of the correct protein construct in the peak fractions was confirmed with SDS-PAGE (e.g. Figure S8B) and Western blot, both demonstrating the presence of different oligomeric protein species in the samples. In all cases, the first peak (INpro-xR-Peak1) eluted close to the void volume of the gel filtration column, indicating the presence of oligomers larger than 700 kDa. This is consistent with the results of Schmid et al. (1997) who showed that the purified INpro molecules had oligomerized ^28^. The additional peaks appeared after the void volume, indicating the presence of smaller oligomers. Using negative stain TEM, we confirmed that the INpro-xR-Peak1 structures are produced by a highly oligomerized form of the protein, substantially larger than the dimensions predicted by our INpro dimer modeling. INpro-67R-Peak1 structures are shown in Figure 4D and E. These structures exhibit an elongated rubber-band-like appearance with a double width of ∼6 nm (∼12 nm in total), which fits well with the dimensions of our modeled INpro dimer and is in good agreement with our cuboid model (Figure 2E, F and H). Overall, the SEC together with the TEM analysis provided experimental support for the elongated shape of INpro and for the higher-order oligomers formed from INpro dimers.

### 2.4. A putative role of C-terminal domain in oligomerization

The role of the N-terminal and C-terminal domains for the ice nucleation activity was investigated with WISDOM using four additional protein constructs, i. e. INpro-16R-ΔN, INpro-15R-ΔT, INpro-N-1R, INpro-C (Figure S3). We observed that the ice nucleation activity of INpro-16R-ΔN was similar to that of INpro-16R that contained an intact N-terminal (Figure S9), indicating that the N-terminal is not necessary for dimer formation and thus, does not affect ice nucleation activity of INpro in general. When both terminal domains were removed (INpro-15R-ΔT, 25 kDa), only a monodisperse peak in SEC was observed, with an elution volume roughly corresponding to the molecular weight of the INpro-15R-ΔT monomer based on calibration of the SEC column using globular standard proteins. A monodisperse peak was previously observed in SEC for a similar INpro-15R-ΔT construct, as well as for other INpro constructs that had their terminals removed ^53^. In addition, no ice-nucleation activity was observed for INpro-15R-ΔT (Figure S9). Based on these results, we conclude that the characteristic nucleation temperature is determined by the CRD size, while the activity depends on the presence of the C-terminal that is involved in INpro oligomerization. It was previously shown that removing the C-terminal from INpro compromised ice nucleation above -5°C ^54^. Han *et al*. , who purified a similar INpro-15R-ΔT, reported that the protein was purified as a monomer and not as a dimer, which would drastically reduce its ice nucleation activity according to the cuboid ice-nucleation model (Figure 2) ^53^. Our conclusion that the C-terminal of the protein plays a role in protein oligomerization is supported by the observation that the C-terminal itself shows weak but significant ice nucleation activity that is constant and independent of the protein concentration. In contrast, the N-terminal itself shows no ice nucleation activity (Figure S9). Cascajo-Castresana et al. recently demonstrated that unspecific protein aggregates can nucleate ice ^55^. We thus suggest that the small but significant degree of ice nucleation activity by the Cterminal could be associated with the C-terminal domains ability to oligomerize into a larger supramolecular structure, which results in a second ice-nucleation mechanism (active at low subzero tempertures) that is distinct from ice-nucleation initiated by the CRD alone but which may be critical for the INpro assembly into dimers and higher order oligo-mers.

## 3. Conclusions

We present the first *ab initio* model of the INpro 3-D structure obtained with AlphaFold and supported by synchrotron radiation circular dichroism data. The proposed model consists of repeat units of two interacting β-strands connected by two sharp turns. Stacking of repeat units results in a β-helix structure with and unusual polar core containing two rows of serine ladders. The two β-sheet faces of the β-helix structure are decorated with the suggested TxT and SxL[T/I] ice-nucleating motifs, respectively. Experimental data on ice nucleation activity of purified INpro constructs were fitted with models based on CNT, which assumed a cuboid shape of INpro derived from the *ab initio* structural model. We thus found that INpro must form dimers to demonstrate observed class C activity. By performing computational docking based on rigid-body algorithms we propose that a parallel dimer structure of INpro forms along the highly conserved tyrosine ladder in CRD yielding a surface where the TxT motif of one monomer aligns with the SxL[T/I] motif on the other. Using transmission electron microscopy, we show the formation of higher-order filamentous INpro oligomers, which we suggest are associated with class A activity. We show that class A activity is maintained with purified protein in the absence of cells and membrane lipids, indicating that higher order oligomers self-assemble from INpro dimers. As dilution of class A INpro leads to the appearance of class C INpro, as observed by LINA and WISDOM measurements, we suggest that higher-level oligomers form through reversible concentration-dependent self-assembly of dimers. Finally, studies using a C-terminal deficient version of the INpro, allows us to conclude that while the characteristic nucleation temperature depends on the number of amino acid repeats in the CRD, its activity ultimately depends on the presence of the C-terminal that seems to be involved in INpro oligomerization. Overall, this study unravels the role of bacterial INpro shape, size and specific oligomerization state for their ice-nucleation activity and develops a theoretical framework of its ice-nucleation activity. Thus, our results form a basis for (i) obtaining a fundamental understanding of ice-nucleation activity in microbial cells in general; (ii) promoting molecular dynamics simulations by providing a testbed and thus bridging the gap between simulation results and experimental data; (iii) the quantitative understanding of the role of INpro in atmospheric ice formation, by providing an opportunity to adopt a theoretical description of INpro for weather and climate modelling; (iv) advocating commercial manufacturing of ice-nucleating particles including INpro class A for industrial applications such as food preservation and crop protection.

## 4. Materials and methods

### 4.1. Modelling of initial 16 repeats of the INpro CRD mono- and dimer

The sequence of the first 16 repeats of the INpro CRD domain was used for ab-initio structure prediction using AlphaFold with a ML algorithm ^56^. The computations were done on the EMCC high performance computing cluster at the Department of Molecular Biology and Genetics at Aarhus University. The structure prediction was performed with all 5 available models and the predicted TM scoring enabled. The best scoring prediction is presented here, and used for the subsequent docking effort. It was selected based on the scoring by the algorithm, by inspection of the predicted aligned error plots, and by visually inspecting the resulting models.

The modelling of the homodimer was performed using the HADDOCK 2.4 webserver ^57^. The tyrosine ladder on both monomers were marked as ‘active residues’. The docking was run with 50,000 structures for rigid body docking, 400 structures for semi-flexible refinement, and 400 structures for final refinement and analysis. All other settings were left as standard. A separate run was performed without marking the tyrosine ladder as active residues. It also showed the tyrosine ladder to be the dimerization interface (data not shown). All figures of the modelling and measurements of the dimensions were made in The PyMOL Molecular Graphics System (Version 2.0 Schrödinger). The sequence identity plot of the INpro CRD was made using Geneious Prime^58^.

### 4.2. Design and Cloning of truncated INpro molecules

As the repeat sequences in the different protein constructs display some sequence variation originating from the naturally occurring repeat sequence divergence in the full length INpro, we calculated the mean similarity scores for the repeats when comparing them to the consensus sequence “GYGSTxTAxxxSxL[T/I]A” ^25^. The similarity scores were assigned to each amino acid in the repeat according to their biochemical properties ^59^. The mean similarity score of single repeats in the CRD ranged from 91.4% to 100% with highly conserved repeats close to the N-terminal and less conserved repeats close to the C-terminal. The difference between the mean similarity score for the different constructs was small and ranged from 94.4% to 98.4% (Figure S4). Based on the low deviation in mean similarity score, we conclude that the repeat sequences are highly similar and show negligible divergence for all constructs.

DNA from strain *Pseudomonas syringae* R10.79 ^41^ was isolated with PowerSoil DNA Isolation kit (Mobio) according to the manufacturer’s instructions. The products were separated on an 1.5% agarose gel and purified using the Nucleotide removal kit (Qiagen). The *ina* gene sequence was obtained from and an *E. coli* codon-optimized synthetic *ina* gene ligated into vector pET24b (Novagen) produced by Genscript ^17^.

Based on the DNA extracted from *P. syringae* R10.79 or the synthetic *ina* gene, we produced 7 truncated versions of the gene encoding for 7 INpro constructs (Tables S1 and S2, Figure S3). The INpro-9R, INpro-16R and INpro-28R contained both terminal domains and a reduced number of amino acid tandem repeats, i.e. 9, 16 and 28 repeats respectively. The The INpro-16R-DN contained only the C-terminal domain and 16 amino acid tandem repeats. The INpro-15R-DT contained only 15 amino acid tandem repeats without the two terminal domains. The INpro-1R-N contained one amino acid tandem repeat with the N-terminal domain and the INpro-C contained just the C-terminal domain. Overlap extension polymerase chain reaction (SOE-PCR) was utilized to obtain genes encoding the INpro-9R, INpro-16R, INpro-16R-DN and INpro-28R as previously described in detail for the INpro-16R ^17^. A regular PCR was used to obtain genes encoding the INpro-15R-DT, INpro-1R-N and INpro-C. All primers are listed in Table S1.

The PCR products were inserted into the pET-30 Ek/LIC vector according to manufacturer’s instructions (LIC Kit, Novagen) (Table S2). The plasmid DNA was isolated using the Gene Jet Plasmid Miniprep Kit (Thermo Scientific) and its sequence was validated by Sanger sequencing (GATC-biotech).

### 4.3. Large scale INpro production

Vectors containing genes encoding for the different protein constructs (Table S2) were transformed into Rosetta (DE3) cells. A single colony of transformed *E. coli* was used to inoculate 50 mL of LB medium with 30 μg/ml Kanamycin and cultured at 37°C overnight. The culture grown overnight was inoculated into 12 L lysogeny broth (LB) medium containing 30 μg/ml Kanamycin for the large-scale expression. Once the culture reached an optical density of 0.6 - 0.8 at 600 nm (OD_600_), it was induced with 1 mM isopropyl β-D-1-thiogalactopyranoside (IPTG) at 20°C overnight. Bacterial cells were harvested by centrifugation at 6,000 × g for 15 min at 4°C. Cell pellets were suspended in 25 ml LB per liter of harvested culture and centrifuged at 4,690 × g for 30 min. Finally, cell pellets were flash-frozen in liquid nitrogen and stored at -80 °C until further treatment.

### 4.4. Protein purification

To purify INpro-9R, INpro-16R, INpro-16R-ΔN, INpro-15R-ΔT, INpro-28R, INpro-1R-N and INpro-C, cells pellet was resuspended in 2 ml of lysis buffer (20mM sodium phosphate, 0.5 M NaCl, 20 mM imidazole; pH 7.4 with a protease inhibitor tablet (Roche) per 1 g of wet cell pellets. Cells were lysed either by 2 rounds of sonication on ice for 5 min (3 cycles at power setting between 70-75%) or with a C5 high-pressure homogenizer (Avestin). During expression tests of INpro-9R, INpro-16R, INpro-16R-ΔN, INpro-15R-ΔT, INpro-1R-N and INpro-C, the proteins were found in both the soluble and insoluble fraction of the cell lysates. For convenience, subsequent work was based on INpro purified from the soluble fractions. Thus, the cell debris and unbroken cells were removed by centrifugation at 12,440 × g for 30 min at 4°C and supernatant was filtered through a 0.45 μm filter. In contrast, INpro-28R was only found in the insoluble fraction. Thus, the cell lysate was spun down for 15 min at 1000 × g, for another 15 min at 15000×g and finally at 42,000 rpm for 2 hours. The membrane was scraped off and resuspended in solubilization buffer (50mM Tris HCl pH 8.0, 300mM NaCl, 5mM MgCl_2_, 30% glycerol) using a glass homogenizer.

The solution containing the proteins was loaded on a prepacked 1 ml or 5 ml His Trap HP Nickel column (GE Healthcare), which had been washed with 5 column volumes of water and equilibrated with 5 column volumes of buffer A (20 mM sodium phosphate, 0.5 M NaCl, 20 mM imidazole; pH 7.4). After washing the column with 10 column volumes of buffer A, the protein sample was eluted on an ÄKTA Start chromatography system with a gradient from 0–100% buffer B (20 mM sodium phosphate, 0.5 M NaCl, 500 mM imidazole; pH 7.4) over 10 column volumes at a flow-rate of 5 ml/min and collected in 5 ml fractions. The protein content of the load, the flow through and fractions of elutes were analyzed on a 12% polyacrylamide SDS-gel. Fractions containing INpro were pooled and for quantification the absorbance was determined at 280 nm using a NanoDrop spectrophotometer. The protein was then concentrated to a final volume of 500 μl using Vivaspin 20 with a molecular weight cut off (MWCO) of 5 kDa (GE Healthcare). The sample was loaded on a Superose6 Increase 10/300 GL size exclusion column from GE Healthcare that had been equilibrated with size exclusion buffer (for INpro-28R: 20 mM Tris-HCl, 150 mM NaCl; pH 7.6, 0.05% n-Dodecyl β-D-maltoside (DDM); and for INpro-9R, INpro16R, INpro-16R-DN, INpro-15R-DT, INpro-1R-N and INpro-C: 20 mM Tris-HCl, 150 mM NaCl; pH 7.6). See an example chromatogram in Figure S8. Fractions of 0.5 ml were collected at a flow of 0.2 to 0.4 ml/min and the protein content was analyzed using a 12% polyacrylamide SDS-gel. Fractions containing the INpro were pooled, frozen in liquid nitrogen and stored at -80°C.

The INpro-67R, which was only found in the insoluble fraction, was extracted and purified using a procedure adapted from Schmid et al. (1997) ^28^. The harvested cells were suspended in 400 ml of cell resuspension buffer (50 mM Tris-HCl, pH 8.0, 100 mM NaCl, 1 mM EDTA, 1 mM 1,4-dithio-dl-threitol, 3mg ribonuclease, 3mg deoxyribonuclease, 5mM benzamidine, 1mM phenylmethylsulphonyl fluoride and a protease inhibitor tablet (Roche)) and then disrupted by a Emulsiflex C5 high-pressure homogenizer (Avestin). Intact bacteria and crude membrane pellets were removed by ultracentrifugation at 144 000 × *g* for 1h at 4°C and the supernatant was diluted 1:1 with 0.2 micrometre filtered milliQ water. An anion-exchange column, HiTrap Q HP column (GE Healthcare), was equilibrated in buffer A (50 mM Tris-HCl pH 8.0, 1 mM EDTA, 1 mM 1,4-dithio-dl-threitol) before loading the protein sample. The flow-through fraction was collected and further purified by ammonium sulfate precipitation. Saturated ammonium sulfate was gradually added to a final concentration of 20% under constant stirring at 4°C. The protein sample was then incubated for 30 mins at 4°C and the precipitate was collected at 14 000 × *g* for 10 min at 4°C. The pellet was re-suspended at 100 mg wet weight/ml in buffer B (50 mM Tris-HCl pH 8.0, 1 mM EDTA, 1 mM 1,4-dithio-DL-threitol, 0.1% N-lauroylsarcosine (NLS)) and incubated for 2 hours at 25°C. The sample was then centrifuged at 10 000 × *g* for 5 min at 20°C to eliminate undissolved pellet particles. The protein sample was then concentrated to a final volume of 500 μl using Vivaspin20 with a molecular weight cut off (MWCO) of 30 kDa. The final protein purification step and determination of oligomeric state was done on a Superose6 Increase 10/300 size exclusion column (GE Healthcare). The column was equilibrated in buffer B and the concentrated protein was loaded onto the column. Fractions of 0.5ml were collected and analyzed using SDS-PAGE. Fractions containing the INpro were pooled, frozen in liquid nitrogen and stored at -80°C.

An overview of the protein purification methods used for all constructs is given in Table S2.

### 4.5. Validation of INpro expression and purification

INpro expression in *E. coli* cells and the presence of purified INpro in fractions from chromatographic methods was confirmed by SDS-PAGE analysis (e.g. Figure S8). Western blot analysis was performed using a primary anti-INpro-1205 rabbit polyclonal antibody, which targets the residues 1205-1220 (CMAGDQSRLTAGKNS) custom-made by Genscript. The blots were probed with a goat anti rabbit secondary antibody (Rockland, USA) (1:5000). Also, Western blots using Anti-His primary antibody (Clontech) (1:4000 dilution) was performed with an anti-mouse HRP conjugate secondary antibody (Dako) (1:3000) to validate the presence of the protein construct.

### 4.6. Synchrotron radiation circular dichroism experiments

SRCD measurements were carried out using the AU-CD beam line of the ASTRID2 synchrotron light source at Aarhus University, Denmark ^60, 61^. INpro-9R and INpro16R samples were prepared in a 20mM NaPhos pH 7.5, 150 mM NaF buffer and spectra measured using a nominally 0.1 mm path length cell (quartz Suprasil cell, Hellma GmbH & Co., Germany), with the precise length determined using interferometry measurements ^62^. The concentration of the samples were determined from the absorbance at 205 nm ^63^, which is measured simultaneously with the CD spectrum. Spectra were measured at 25°C in triplicate in the wavelength range of 170 to 280 nm in steps of 1 nm and a dwell time of 2 secs, with a corresponding baseline measured on the buffer alone. SRCD data were deconvoluted using the CDSSTR analysis programme and SMP180 reference datasets at the DichroWeb portal ^42, 43^. The secondary structure content for the 7 repeats representing the difference between the INpro-9R and INpro16R constructs was calculated based on the SRCD data given the cumulative nature of SRCD data.

### 4.7. Transmission Electron Microscopy

All EM work was done at the iNANO EM facility (EMBION), Aarhus University. Copper grids with a 400 square mesh were prepared with a 2% Collodion solution and carbon coated using a Leica EM SCD 500. The grids were freshly glow-discharged with a PELCO easiGlow™ Glow Discharge Cleaning System prior to loading them with 5µL of protein sample, blotting and staining 3-times with 3 µL 2% uranyl formate. Samples dilutions 1:5, 1:10, 1:100 and 1:1000 were applied to the grids. TEM images were acquired at a nominal magnification of 67,000× (pixel size size of 1.67 Å) using a FEI Tecnai Spirit transmission electron microscope with a TWIN lens operating at 120 kV. Images were collected with a Tvips TemCam F416 CMOS camera.

### 4.8. WISDOM setup and sample handling

Ice nucleation measurements were conducted using the WeIzmann Supercooled Droplets Observation on a Microarray (WISDOM). The setup, validation process and temperature calibration are discussed in details previously ^44, 64, 65^ thus only a general description is given here.

We used a microfluidic device to suspend droplets (with a diameter of 100 µm) containing the proteins in an oil mixture (2% wt Span 80 in mineral oil). Droplet generation was done using NE-500 Programmable OEM Syringe Pump to inject the oil and the sample simultaneously into the device in separate inlets, allowing the oil to press and cut monodisperse droplets in a four-way junction. Droplet size was controlled by changing the ratio of the sample to oil flow. The droplets’ diameter in this study was about 100 µm. Droplets are then trapped in the device’s chamber array, transferred immediately to a Linkam cooling stage (LTS420) coupled to an optical microscope (Olympus BX51) and a CCD camera. The temperature ramp in this study included three different cooling rates: first, droplets were cooled fast at 20℃ min^-1^, from the room temperature, in order to reach fast to the region where the proteins were ice-active. Then, a slower cooling rate of 5℃ min^-1^ was applied to reach closer to the onset of the freezing, and finally, slow cooling rate of 1℃ min^-1^ to follow accurately the freezing events . Freezing events were detected optically, at the point where the droplet becomes darker due to the light scattering of the ice crystals. After all the droplets were frozen, they were heated fast until the melting point, that was recorded at a rate of 1℃ min^-1^.

To retrieve the INpro spectra for a wider range of temperatures, samples were analyzed at several concentrations by serially diluting the sample after each analysis by a factor of 10 or 100. The dilution was performed with the buffer until homogenous freezing was observed. Analysis was repeated at least three times for each concentration for statistical validation. The buffer of the INpro-28R and the INpro-67R contained some detergent that interacted with the WISDOM’s oil so we could not produce drops using the buffer. To overcome this issue, we diluted these samples with deionized water by a factor 5 and 2.5 for the INpro-28R and INpro-67R, respectively. Figures 2B and C present a scheme of the WISDOM setup.

### 4.9. LINA setup and sample handling

In order to analyze INpro that are present at low concentration, we performed complementary experiments with a second droplet freezing array LINA (Leipzig Ice Nucleation Array) ^48^, based on a previously described instrument ^30^. LINA measurements were carried out with 90 droplets each having a volume of 1 µl. These droplets are around 5x10^5^ larger than droplet volumes used for WISDOM, allowing for the detection of rarer INpro. Droplets were placed in separate compartments on a hydrophobic glass slide on top of a cooled Peltier element. We used a standard cooling rate of 1 K/min, recorded the freezing droplets optically and obtained a temperature resolution of 0.1 K with an uncertainty below 0.18 K (single standard deviation) ^66^.

Similar as WISDOM measurements, we made droplet freezing experiments of dilution series of the INpro samples with the respective buffer to retrieve INpro spectra for a wide range of temperatures until we observed a levelling off indicating that all existing INpro were already activated.

### 4.10. The characteristic nucleation temperature (T_char,50_)

In general, the freezing probability of a droplet population depends on the properties and number of immersed ice nucleating particles (INP). Consequently, within certain limits determined by the INP properties, the higher the number of immersed INP the higher is the temperature range at which the freezing is observed ^67–69^. In order to define a characteristic nucleation temperature, independently of the number of INP, which in our case are INpro, the related calculations are done based on the ice nucleation number site density *N*_*n*_. In some cases, the ice nucleation spectrum exhibits a clear plateau ^67, 69, 70^. This is indicative of the fact that there are no further INP in the sample which nucleate ice at the respective temperatures for which the plateau shows up. Hence from this, the maximum number of INP of the respective freezing mode can be derived. In this study, instead of observing a clear plateau, we observed that *N*_*n*_ continued to slightly increase with decreasing temperature. We assume this to be due to eventually existing degradation products of INpro constructs, which are less efficient in nucleating ice. However, as we do not have such a clear plateau, we determined the knee point of the *N*_*n*_ spectra, which we assume to be the maximum number of INpro. This was done by using python function kneed which uses temperature bin averaged logarithmic ice nucleation spectra as input data (Figure S7, Table S3). Based on this knee point, the characteristic nucleation temperature (T_char,50_) is defined as the temperature at which the value of *N*_*n*_ is 50% of *N*_*n*_ at the knee point. The *N*_*n*_ at the knee point ranged roughly from 10^-10^ to 10^-5^ ice-nucleation active INpro molecules per total number of INpro molecules.

### 4.11. Theoretical description of elongated ice cluster templates using Classical Nucleation Theory

Considering heterogeneous ice nucleation, the most commonly used geometry of ice clusters on a flat surface is the spherical cap in CNT ^46, 71^. Ice nucleating particles providing these surfaces for ice cluster formation might have special features that require other geometries, such as hexagonal shape, curved surfaces, pores, completely covered or elongated templates to be described ^29, 72–75^.

As the shape of the CRD region of the analyzed proteins is much more elongated than wide, we consider these elongated templates for ice nucleation. We assume that the relevant surface area for ice nucleation is characterized by a long and a short axis. We model a simple cuboidal ice cluster which forms at the surface of the elongated ice cluster template (Figure 2E and F). The cuboid has three axes: *a* represents the longest axis, *b* the width and *c* the height.

To form an ice cluster (2, ice) in a metastable parent phase (1, supercooled water) on a template (3, protein structure) energy is released by forming a volume *V* and consumed to form a surface at the *ij* and a line at the *ijk* interfaces. The Gibbs free energy difference between the liquid and the ice phase Δ*G* assuming the cuboidal geometry of the ice cluster can be written as:

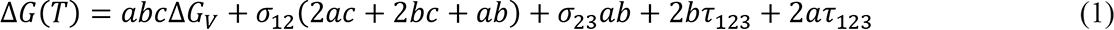

with the respective interfacial free energies σ_*ij*_, line tensions τ_*ijk*_ and Δ*G*_*V*_(−*kT*/*v*_*ice*_(*T*)*ln*(*S*_12_) including with Boltzmann’s constant *k* = 1.38065 × 10^−23^m^2^kgs^-2^K^-1^, the ratio of saturation pressure of liquid water to ice *S*_12_ and the molecular volume of ice *v*_*ice*_ (parameterizations used are given in ^76, 77^).

As the width of the template and therefore the width of the ice cluster *b* is limited by the properties of the INpro for this special geometry, we can justify the assumption *b* = *const*. In order to derive the critical size of the ice cluster, Eq. (1) is differentiated with respect to c and set to zero. We obtain the critical length *a*_*crit*_ with

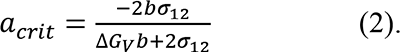

Thus, we obtain the critical height of the cuboidal ice cluster. As a result, the critical length *a*_*crit*_ and height (not shown) are independent of each other, but depend both on the constant width *b* of the cuboid. In the framework of the cuboidal model we consider the critical length *a*_*crit*_ as characteristic length as it defines basically the interface area between ice cluster and template.

In the model using the spherical cap geometry (Figure 2G), the ice cluster can be described by the critical radius 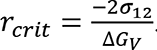. Following this approach, the radius of the interface circular area between ice cluster and nucleation template *r* = *r*_*crit*_ × *sinθ* determines the characteristic length (Figure 2G).

In Figure 2H the characteristic lengths of cuboid and spherical cap model as functions of temperature are shown. In general, the higher the temperature the larger is the characteristic length needed to form critical ice clusters in both models. Further the characteristic lengths derived from the cuboidal model are always larger, i.e., energetically less efficient, compared to the spherical cap model. For very high widths *b*, *a*_*crit*_ converges to *r*_*crit*_.

At this point we would like to discuss the simplified assumption of a cuboid geometry as a real ice cluster would never take such a shape. Probably a more realistic shape would be an ellipsoidal cap in analogy to the spherical cap or a cuboid with rounded off edges ^29^. However, the relevant characteristic length is expected to be very similar in assuming these geometries. Consequently, we refrain from considering these much more complicated approaches.

## ASSOCIATED CONTENT

### Supporting Information

Figure S1. Additional information from Alphafold for the predicted structure of INpro CRD repeat 1-16.

Figure S2. Alphafold structure prediction of INpro-67R.

Figure S3. Sketch of all protein constructs used for the study.

Figure S4. The mean similarity scores of all repeats for the different INpro constructs.

Figure S5. Synchrotron Radiation Circular Dichroism of INpro-9R and INpro-16R

Figure S6. Ice-nucleation spectra for INpro-9R, INpro-16R, INpro-28R and INpro-67R.

Figure S7. Examples of knee point detection applying the sensitivty parameter 1.0.

Figure S8. Example size exclusion chromatogram and SDS-PAGE gel image.

Figure S9. Frozen fraction as a function of temperature for INpro-16R-DN, INpro-15R-DT samples and the C-Terminal and N-Terminal samples measured by WISDOM.

Table S1. Primers used for different protein constructs.

Table S2. An overview of cloning and purification details for all protein constructs used in the study.

Table S3. Properties of the different protein constructs: presumed state of oligomerization, geometry, INpro class with respective characteristic nucleation temperatures measured with WISDOM and LINA instruments.

## ACKNOWLEDGMENT

Author contributions: K.F., T.B, M.L. and T.Š-T. designed the research project. M.L., T.B. and L.S.A.D. designed the protein constructs and optimized the purification. T.Š.-T. and T.B. supervised the project. M.L., L.Z.J., S.B., N.R. A.Z., S.G, and S.H. performed the experiments. T.D, T.B., N.C.J. and S.V.H. performed SRCD analysis. L.S.A.D. performed the ab initio modeling under supervision of T.B.. S.H. and D.N. performed the CNT modeling. T.Š-T. wrote the manuscript with contributions from all coauthors.

## Data and materials availability

All data needed for evaluating the paper are presented in the main body and in the Supplementary Materials. Additional data are available upon request from the authors.

## Supplementary Materials

Figure S2. Alphafold structure prediction of INpro-67R.

Figure S3. Sketch of all protein constructs used for the study.

Figure S4. The mean similarity scores of all repeats for the different INpro constructs.

Figure S5. Synchrotron Radiation Circular Dichroism of INpro-9R and INpro-16R

Figure S6. Ice-nucleation spectra for INpro-9R, INpro-16R, INpro-28R and INpro-67R.

Figure S7. Examples of knee point detection applying the sensitivty parameter 1.0.

Figure S8. Example size exclusion chromatogram and SDS-PAGE gel image.

Figure S9. Frozen fraction as a function of temperature for INpro-16R-ΔN, INpro-15RΔT samples and the C-Terminal and N-Terminal samples measured by WISDOM.

Table S1. Primers used for different protein constructs.

---

**Figure S1.**
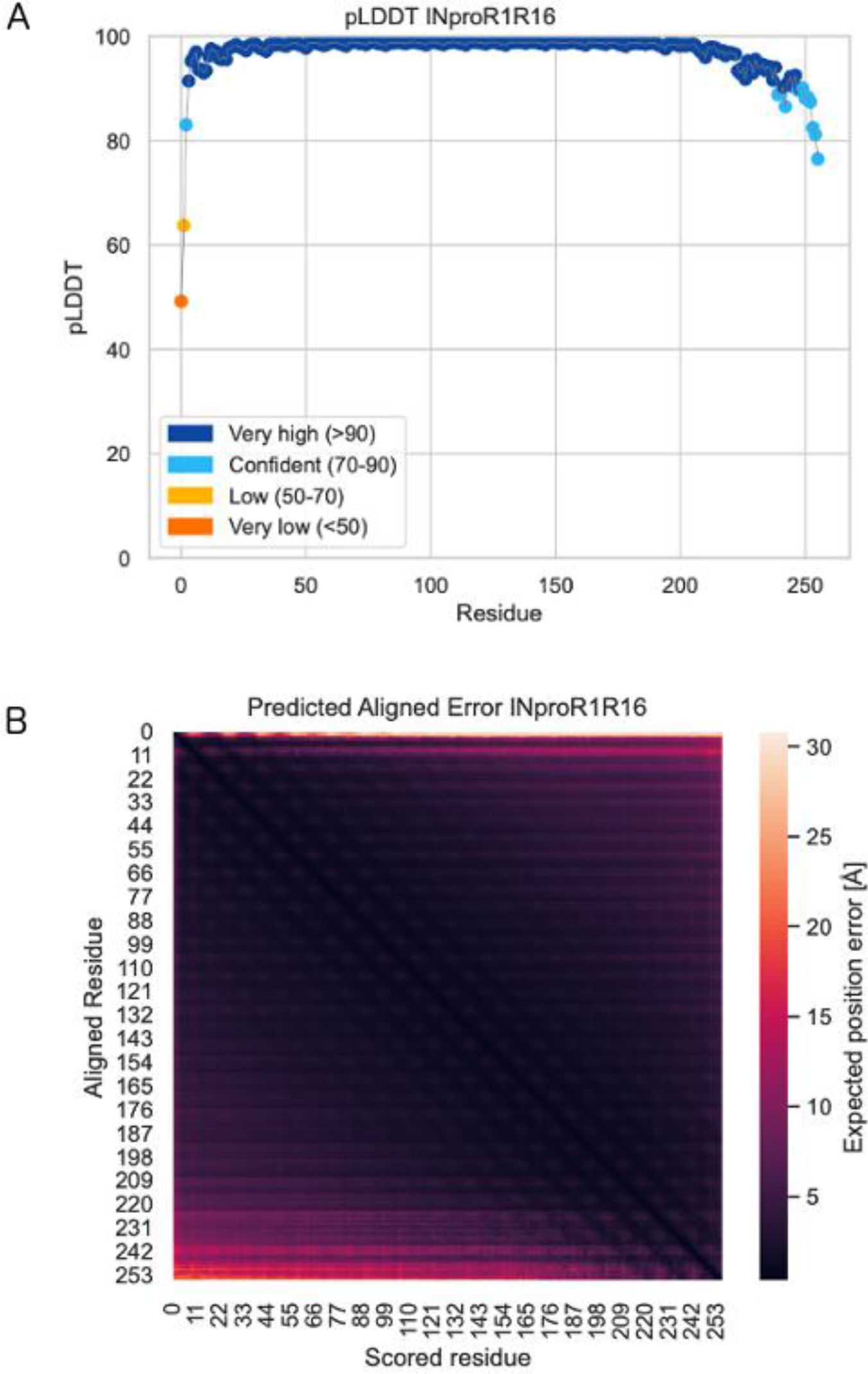
Additional information from Alphafold for the predicted structure of INpro CRD repeat 1 to repeat 16. The structure is presented in the main paper. **(A)** pLDDT score as a function of residue number. The score ranges from 0-100, with higher scores indicating a higher confidence. (B) Predicted aligned error plot. This plot shows the expected position error between pairs of residues. Hereby revealing the confidence in the overall topology of the protein, and the predicted domains. It is shown as residue vs residue. Dark colors show increased confidence.

**Figure S2.**
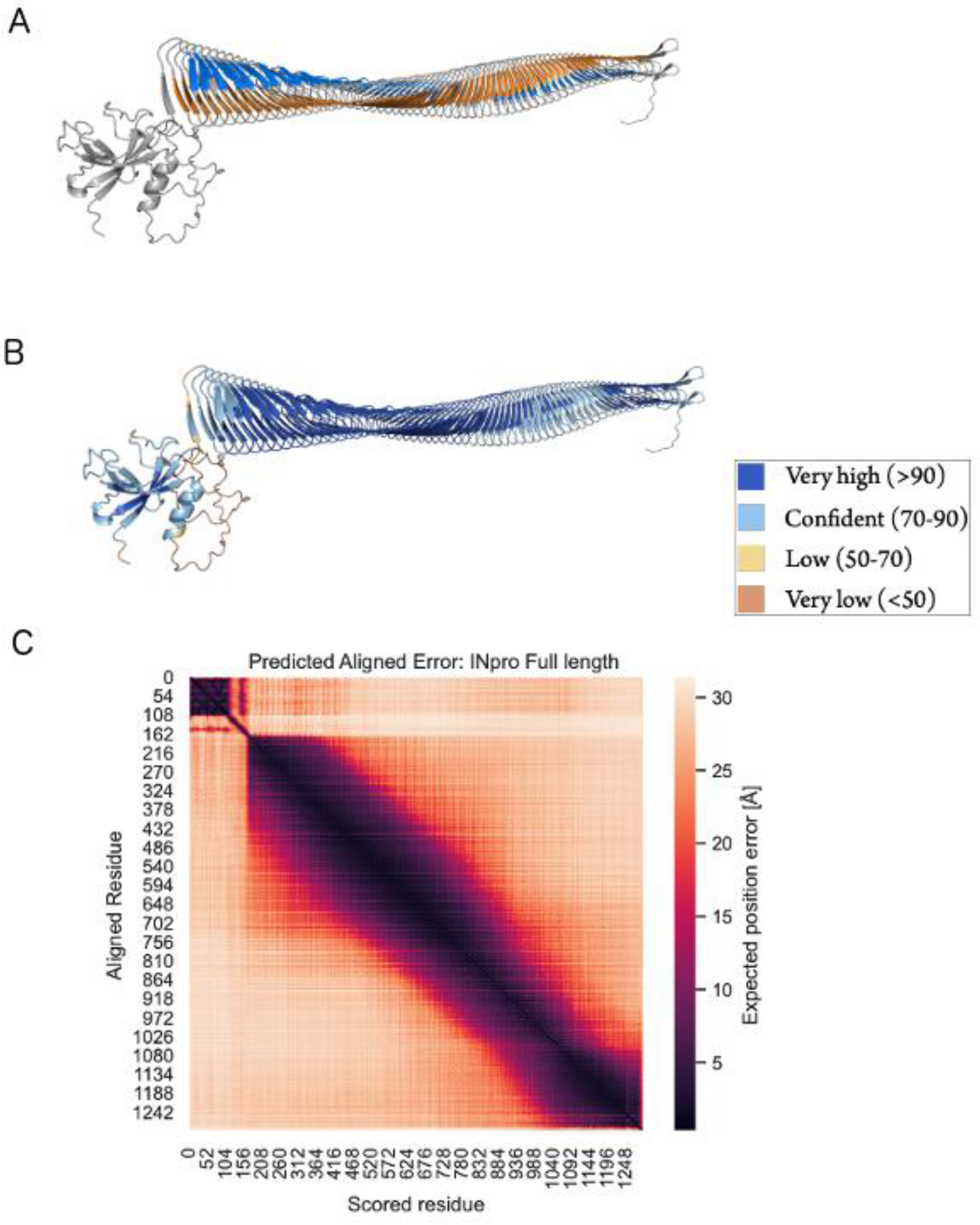
Alphafold structure prediction of INpro-67R. **(A)** The predicted structure colored in accordance with figures in the main paper. Alphafold predicts a folded domain in the N-terminal region (top, left), an unstructured (or of unknown fold) linker, a beta-helix CRD domain, and short capping of the beta-helix by the C-terminal domain ending in an unstructured (or of unknown fold) part. **(B)** Same figure colored according to the Alphafold pLDDT score. The score ranges from 0-100, with higher scores indicating a higher confidence. **(C)** Predicted aligned error plot from Alphafold. This plot shows the expected position error between pairs of residues. Hereby revealing the confidence in the overall topology of the protein, and the predicted domains. It is shown as residue vs residue. Dark colors show increased confidence

**Figure S3.**
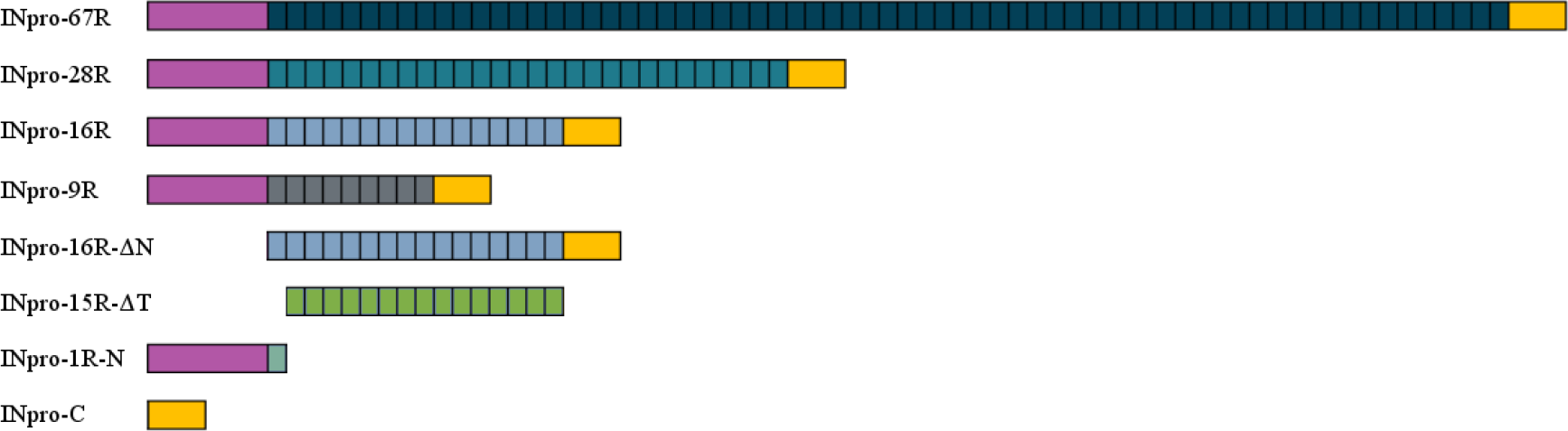
Sketch of all protein constructs used for the study. Purple color signifies the N-terminal domain and the yellow color signifies the C-terminal domain. The blue and green shades in the CRD are symbolic and are used to emphasize the different INpro constructs.

**Figure S4.**
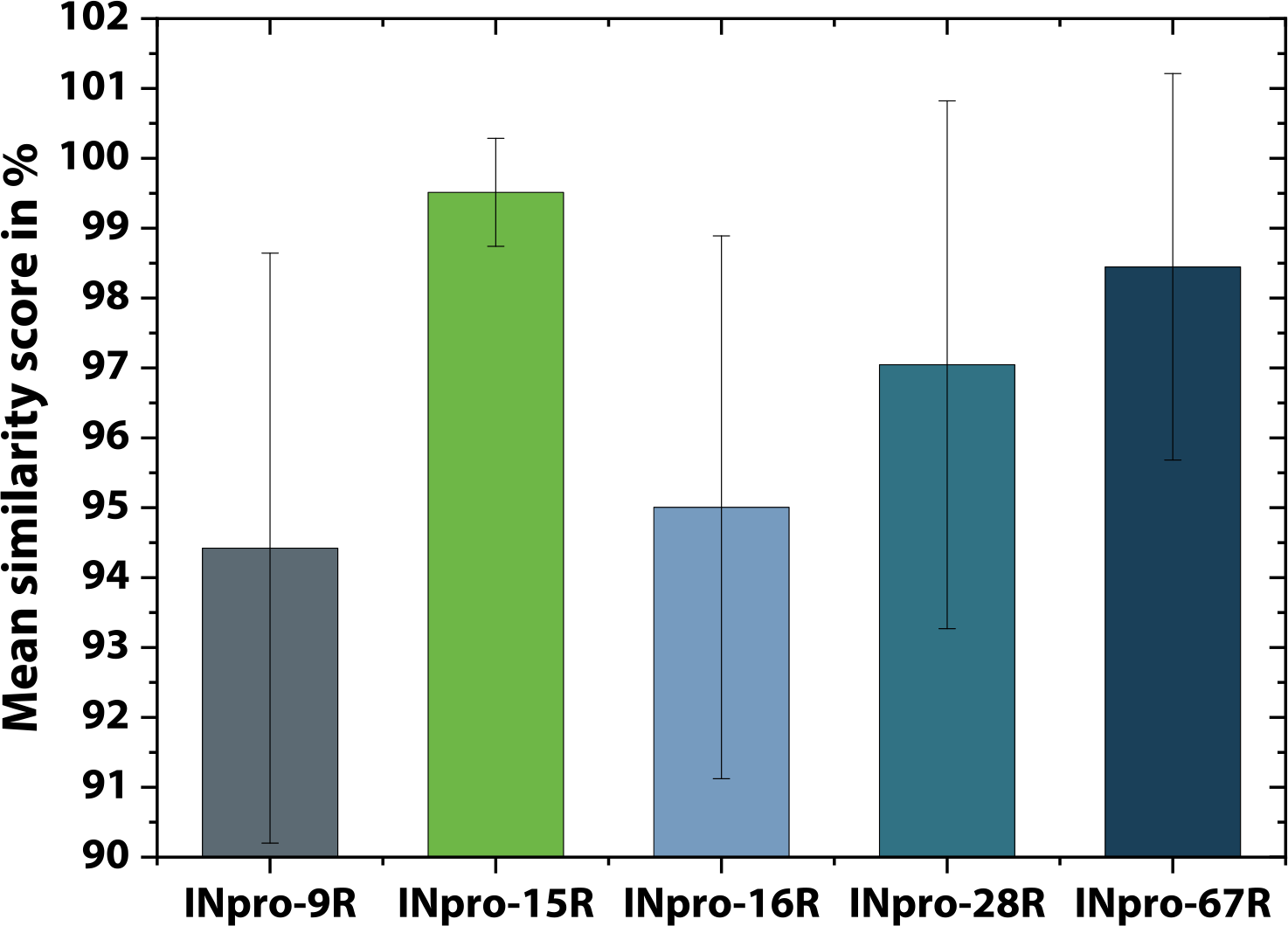
The mean similarity scores of all repeats for the different INpro constructs.

**Figure S5.**
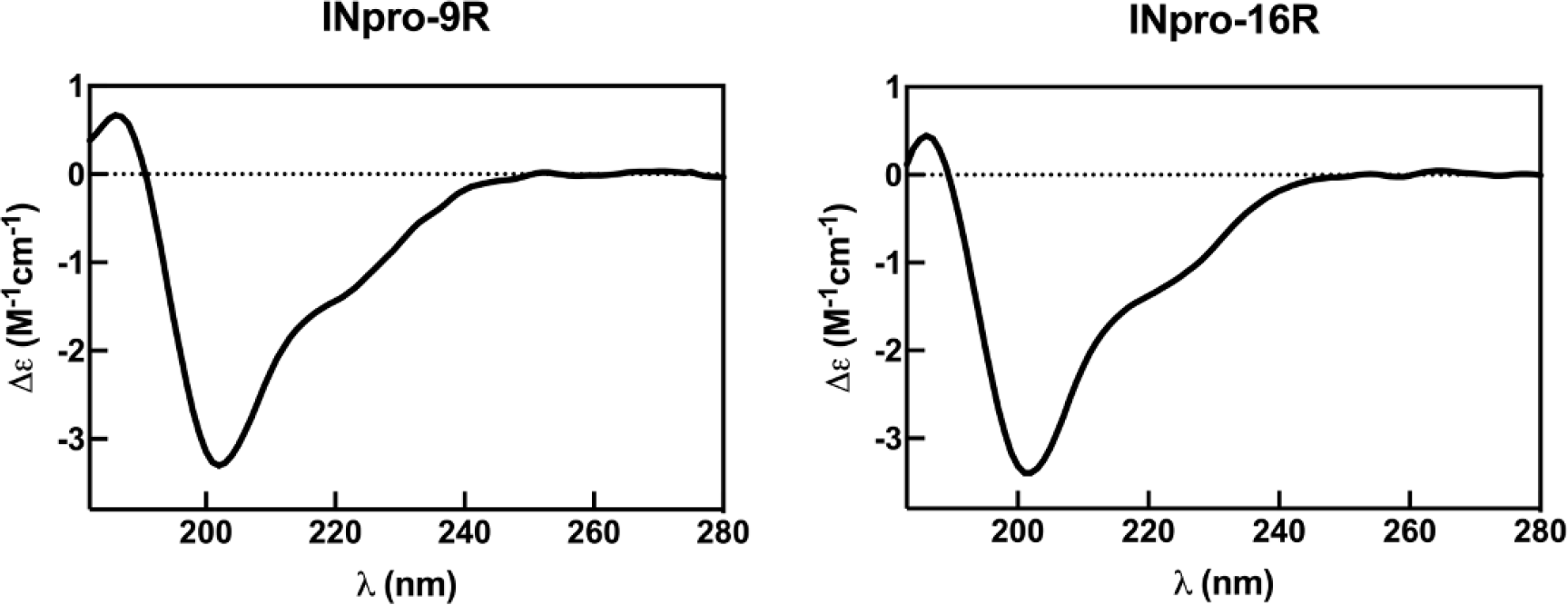
Representative synchrotron radiation circular dichroism spectra for INpro-9R and INpro-16R showing beta strand/coil/turn structure features when deconvoluted with the SMP180 reference datasets at the DichroWeb portal. Data are shown for the wavelength range 182-280 nm.

**Figure S6.**
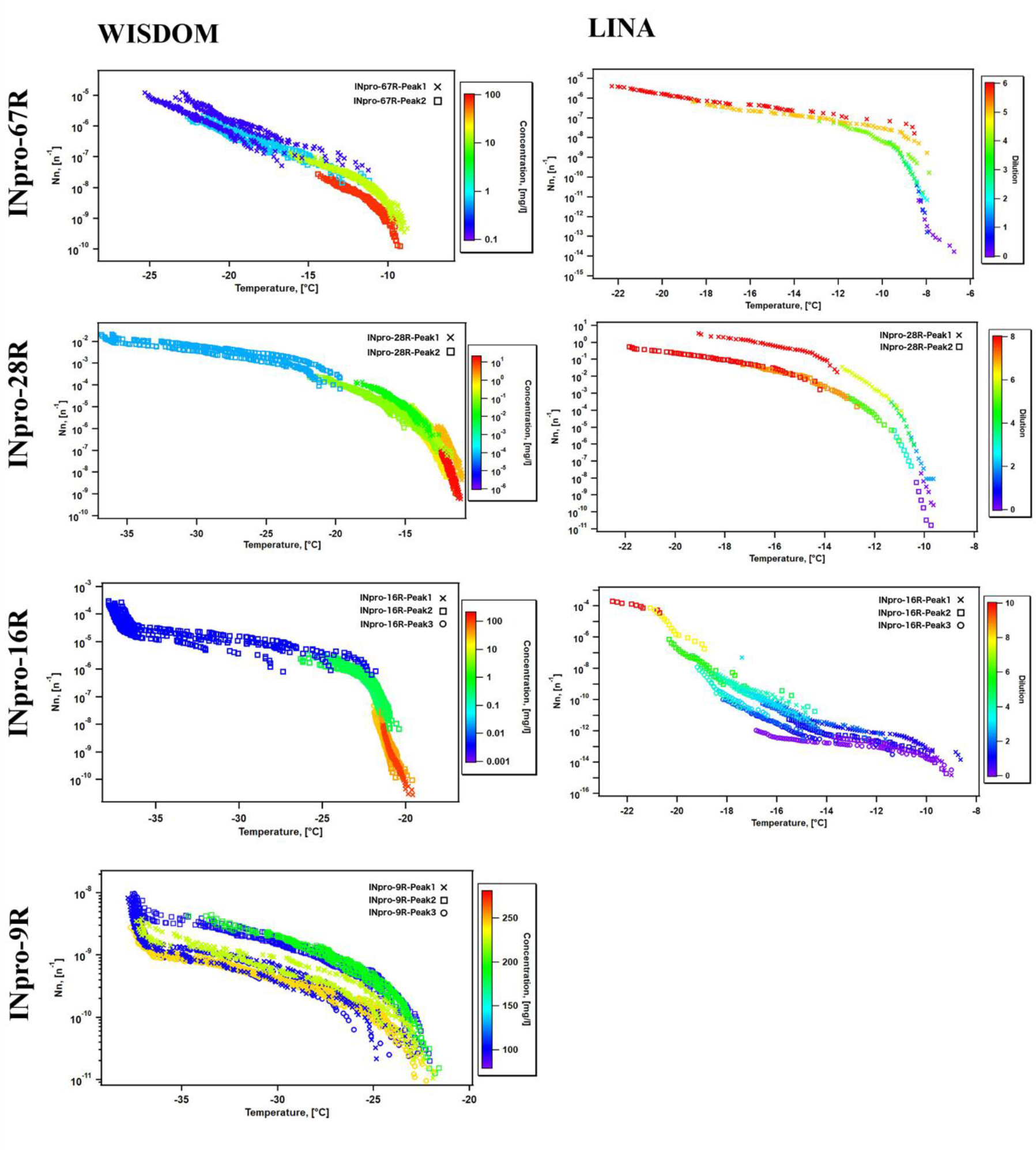
Ice-nucleation spectra for INpro-9R, INpro-16R, INpro-28R and INpro-67R. N_n_ is shown as a function of temperature as measured by WISDOM and LINA. The different symbols represent different peaks of the same sample (P1, P2, P3). Plots on the left are color coded with respect to concentration and plots on the right with respect to dilution (10-fold dilutions are presented on a logarithmic scale). The fact that *N*_*n*_ at the knee point for the different protein constructs ranged over several orders of magnitude (from 10^-10^ to 10^-5^) indicated that the difference in purification procedures (18, 63–66) and the stability of the individual constructs had an effect on the ratio between the correctly folded and active INpro molecules and all INpro molecules (or total purified protein mass as measured by the absorbance at 280 nm) in individual samples. The fact that there is still some increase in N_*n*_ below the steepest region can be explained by the presence of a heterogeneous population of ice nucleation active compounds, which is likely due to the presence of INpro degradation products (63–65). Periodic proteolytic degradation of INpro, which decreased the nucleation temperature, has previously been reported for INpro (63–65).

**Figure S7.**
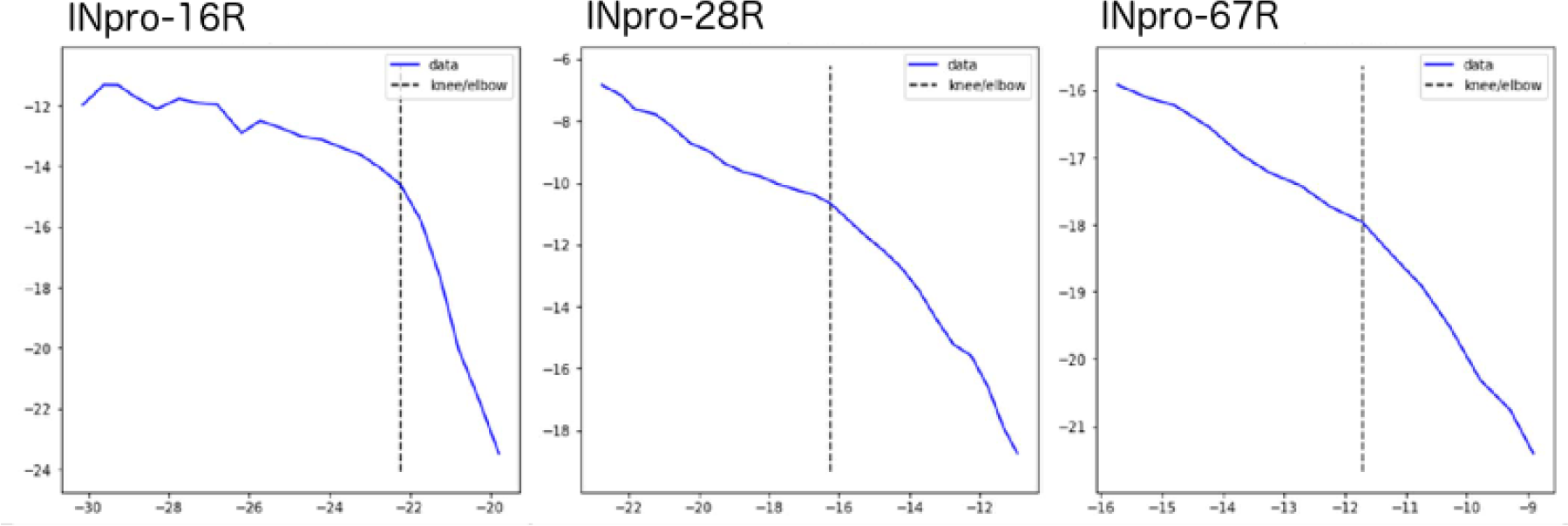
Examples of knee point detection applying the sensitivty parameter 1.0.

**Figure S8.**
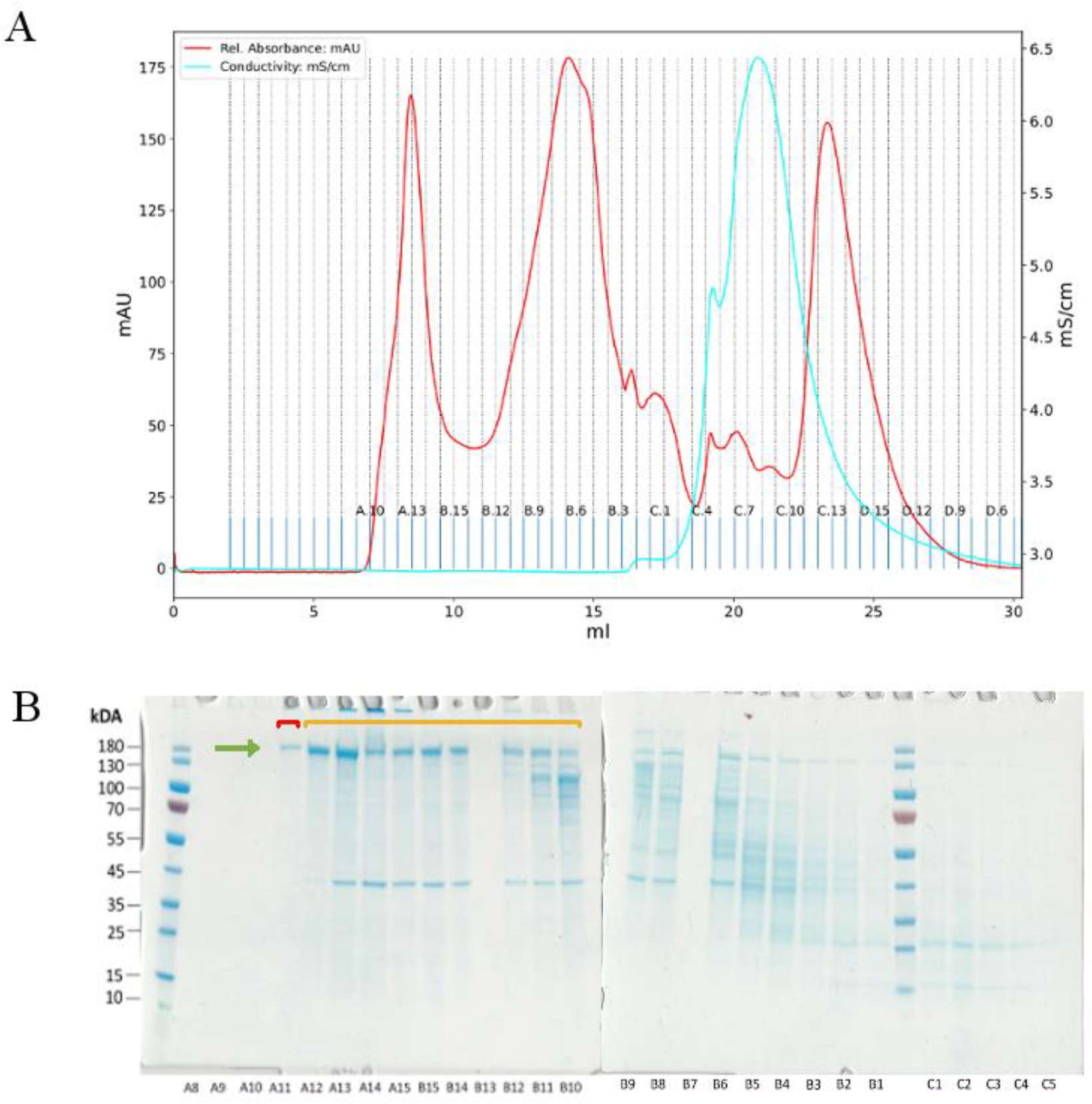
Example size exclusion chromatogram and SDS-PAGE gel image. **(A)** A chromatogram from a size-exclusion chromatography (SEC) of the INpro-67R is shown. Absorbance at 280nm is shown for individual 0.5 mL fractions. **(B)** SDS-PAGE gel for the selected fractions is shown. The green arrow indicates the band corresponding to the INpro-67R. Even though the theoretical molecular weight of INpro-67R was 127.26 kDa the protein migrated just above the 180 kDa marker. Its identity was veryfied by Western blotting (data not shown). The first eluted fraction was stored as INpro-67R-Peak1 (shown in red). The following fractions were pooled as INpro-67R-Peak2 (shown in orange).

**Figure S9.**
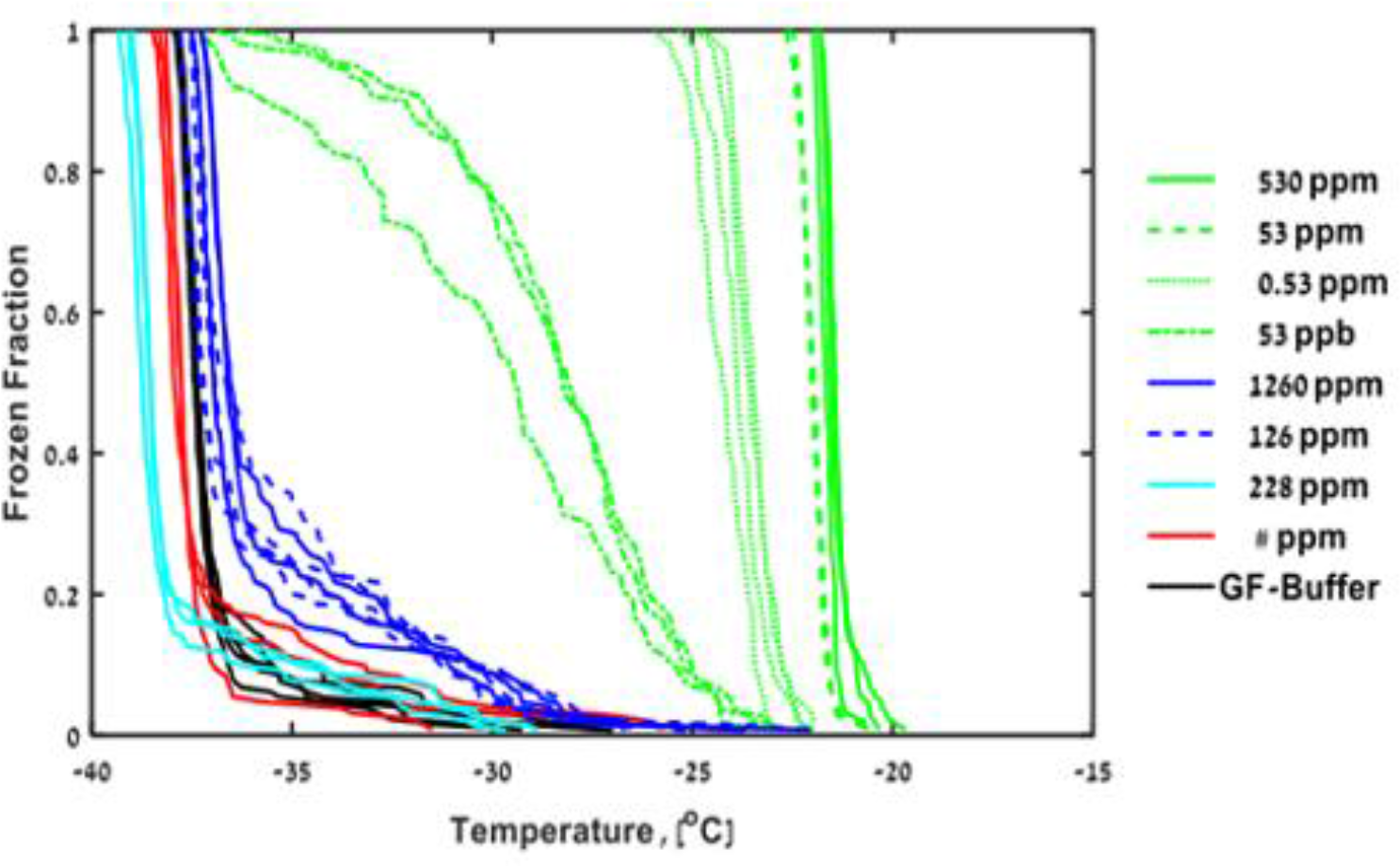
Frozen fraction as a function of temperature for INpro-16R-DN (green), INpro-C (blue), INpro-N-1R (cyan) and INpro-15R-DT (red) samples measured by WISDOM.

**Table S1:**
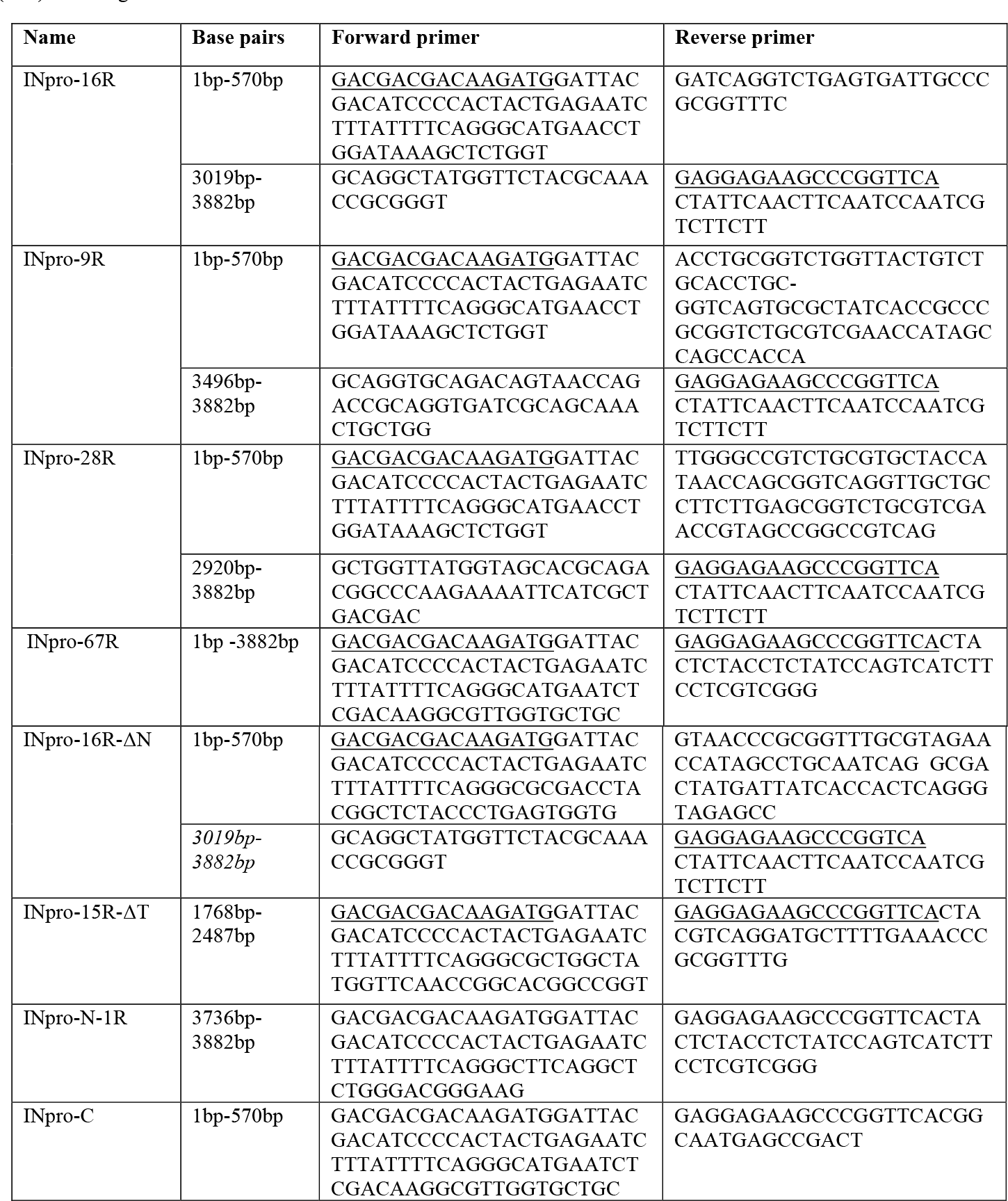
Primers used for different protein constructs. The underlined sequences correspond to the ligation-independent cloning (LIC) overhangs.

**Table S2:** An overview of cloning and purification details for all protein constructs used in the study. *AC – affinity chromatography; SEC – size-exclusion chromatography; AEC – Anion exchange chromatography; ASP – Ammonium sulphate precipitation. All strains were expressed in Rosetta (DE3) Competent E. coli Cells (Novagen).

| Protein construct | Vector | Tag | Tev site | Mol. weight [kDa] | Purification steps* | Detergent in the final sample |
| --- | --- | --- | --- | --- | --- | --- |
| INpro-9R | *EK lic pET30* | N-terminal His Tag and S Tag | Y | 41 | AC, SEC | No detergent |
| INpro-16R | *EK lic pET30* | N-terminal His Tag and S Tag | Y | 53,7 | AC, SEC | No detergent |
| INpro-28R | *EK lic pET30* | N-terminal His Tag and S Tag | Y | 72 | AC, SEC | 0.05% n-Dodecyl β-D-maltoside (DDM) |
| INpro-67R | *EK lic pET24b+* | C-terminal His Tag | N | 143,6 | AEC, ASP, SEC | 0.1% N-lauroylsarcosine (NLS) |
| INpro-16R-ΔN | *EK lic pET-30* | N-terminal His Tag and S Tag | Y | 37,6 | AC, SEC | No detergent |
| INpro-15R-ΔT | *EK lic pET30* | N-terminal His Tag and S Tag | Y | 25 | AC, SEC | No detergent |
| INpro-1R-N | *EK lic pET46* | N-terminal His Tag | Y | 21,6 | AC, AEC, SEC | No detergent |
| INpro-C | *EK lic pET41* | N-terminal GST Tag , S tag and His Tag | Y | 31,5 | AC, AEC, SEC | No detergent |

**Table S3:** Properties of the different protein constructs and oligomerization states. Tchar,knee, Tchar,50, Tchar,10, Tchar,1 are the characteristic nucleation temperature of homogeneous INpro at 100%, 50%, 10% and 1% of the knee point.

| Protein construct | Presumed state of oligomerization | Length /nm<sup>a</sup> | Width /nm<sup>a</sup> | INpro class | $N_{n,knee} /n^{-1}$ | $T_{char,knee}/^{\circ}C$ | $T_{char,50}/^{\circ}C$ | $T_{char,10}/^{\circ}C$ | $T_{char,1}/^{\circ}C$ | instrument |
| --- | --- | --- | --- | --- | --- | --- | --- | --- | --- | --- |
| INpro-9R | dimer | 4.1 | 5.5 | C | $2.55 \times 10^{-10}$ | -24.7 | -23.7 | -22.3 | -21.8 | WISDOM |
| INpro-16R | dimer | 7.3 | 5.5 | C | $4.44 \times 10^{-7}$ | -22.2 | -21.7 | -21.3 | -20.8 | WISDOM |
| INpro-28R | dimer | 12.8 | 5.5 | C | $2.33 \times 10^{-5}$ | -16.3 | -15.8 | -14.2 | -12.8 | WISDOM |
| INpro-67R | dimer | 30.6 | 5.5 | C | $1.59 \times 10^{-8}$ | -11.7 | -11.2 | -9.8 | -8.9 | WISDOM |
| INpro-67R | dimer | 30.6 | 5.5 | C |  |  |  |  |  | LINA |
| INpro-67R | Higher order oligomer | na | na | A |  |  |  |  |  | LINA |
| INpro-16R | dimer | 7.3 | 5.5 | C | $4.44 \times 10^{-7}$ | -22.2 | -21.7 | -21.3 | -20.8 | LINA |
| INpro-16R | Higher order oligomer | na | na | A | $1.25 \times 10^{-14}$<sup>b</sup> | -9.6 | -9.3 | -9.0 | / | LINA |
| *E.coli* expressing INpro-67R | dimer | / | / | C |  |  |  |  |  | LINA |
| *E.coli* expressing INpro-67R | Higher order oligomer | / | / | A | $4.04 \times 10^3$<sup>c</sup> | -4.7 | -4.5 | -4.1 | -3.8 | LINA |

<sup>a</sup> The length and the width are derived from the structural INpro model.
<sup>b</sup> Knee point and derived values have higher uncertainties as the data base is low.
<sup>c</sup> $N_v$ instead of $N_n$

